# Enzymatic dissociation induces transcriptional and proteotype bias in brain cell populations

**DOI:** 10.1101/2020.05.14.095422

**Authors:** Daniele Mattei, Andranik Ivanov, Marc van Oostrum, Stanislav Pantelyushin, Juliet Richetto, Flavia Mueller, Michal Matheus Beffinger, Linda Schellhammer, Johannes vom Berg, Bernd Wollscheid, Dieter Beule, Rosa Chiara Paolicelli, Urs Meyer

## Abstract

Different cell isolation techniques exist for transcriptomic and proteotype profiling of brain cells. Here, we provide a systematic investigation of the influence of different cell isolation protocols on transcriptional and proteotype profiles in mouse brain tissue by taking into account single-cell transcriptomics of brain cells, proteotypes of microglia and astrocytes, and flow cytometric analysis of microglia. We show that standard enzymatic digestion of brain tissue at 37°C induces profound and consistent alterations in the transcriptome and proteotype of neuronal and glial cells, as compared to an optimized mechanical dissociation protocol at 4°C. These findings emphasize the risk of introducing technical biases and biological artefacts when implementing enzymatic digestion-based isolation methods for brain cell analyses.

## Introduction

The neuroscience field is constantly implementing novel techniques for gene sequencing, epigenetic analyses and proteotype profiling at a rapidly increasing rate. Obtaining single cell suspensions from intact brain tissue is essential for downstream cell specific transcriptomic and proteotype analyses. The main techniques available to obtain single cell suspensions from brain tissue are enzymatic digestion (ED) and mechanical dissociation (MD). The latter is typically carried out at 4°C (Mattei et al., 2017), whereas ED requires incubation at temperatures between 30-37°C (Saunders et al., 2018; Tiklova et al., 2019; Zhang et al., 2014). With both techniques, cells such as neurons and microglia will lose their processes and suffer harsh conditions whereby a proportion of cells die during the dissociation process. Although loss of cells and of their processes is likely to introduce technical artefacts in both cases, we hypothesize that at warm temperatures, as opposed to 4°C, surviving cells may overreact to the non-physiological milieu containing shredded extracellular matrix, cell debris, and cytoplasmic leakage (Reichard & Asosingh, 2019). Furthermore, the ED technique is associated with marked temperature shifts occurring between the eventual cell isolation step and the preceding perfusion which is typically carried out at cold temperatures to remove blood contamination from the brain. Thermal shocks resulting from temperature shifts have been shown to elicit widespread transcriptional changes, even within short time windows (Mahat, Salamanca, Duarte, Danko, & Lis, 2016). Hence, unlike MD at cold temperatures, which largely spares the cellular transcriptome and proteome (Fujita, 1999), the ED technique may introduce additional biological biases to the transcriptional and proteotype profiles of cells.

A few studies have thus far warned for possible ED-induced artefacts, but not as a main focus of investigation, and proposed the use of transcriptional/translational blockers to prevent *de novo* transcription/translation (Ayata et al., 2018; Hrvatin et al., 2018; Wu, Pan, Zuo, Li, & Hong, 2017). Nevertheless, this issue has only been considered for transcriptional analyses, whereby possible biological consequences of ED on the proteotype profiling (e.g. flow cytometry and mass-spectrometry analyses) has up to today not been taken into consideration. Importantly, the use of transcription/translation inhibitors do not prevent biological downregulation of RNAs and proteins that might occur during ED, nor can they prevent unwanted alterations in epigenetic marks or internalization/externalization of receptors which might be crucial readouts for certain cell specific analyses. Given that at present, ED is the most widely used method to obtain single cell suspensions from brain tissue (Figure S1), it is pivotal to understand the extent of cell responses when exposed to ED as compared to a method which holds cells in a quiescent state. We here investigate systematically the influence of different cell isolation protocols on transcriptional and proteotype profiles in mouse brain tissue, by taking into account single-cell transcriptomics, proteotypes of microglia and astrocytes, and flow cytometric analysis of microglia. By comparing the transcriptional and proteotype profiles of brain cells isolated via ED at 37°C and via MD at 4°C, our study provides a comprehensive transcriptomic and proteotype resource that can be used to identify and correct for artefacts in previously published data sets generated via ED. Furthermore, with the present work, we share an optimized and easy-to-establish MD protocol at 4°C, which is cost effective and technically easier to implement than conventional ED-based protocols.

## Material and Methods

### Animals

Nine to ten weeks old male C57BL6/N mice (Charles River Laboratories, Sulzfeld, Germany) were used throughout the study. They were caged 3-5 animals per cage in individually ventilated cages (IVCs). The animal vivarium was a specific-pathogen-free (SPF) holding room, which was temperature- and humidity-controlled (21 ±3 °C, 50 ± 10%) and kept under a reversed light–dark cycle (lights off: 09:00 AM–09.00 PM). All animals had ad libitum access to the same food (Kliba 3436, Kaiseraugst, Switzerland) and water throughout the entire study. All procedures described in the present study had been previously approved by the Cantonal Veterinarian’s Office of Zurich, and all efforts were made to minimize the number of animals used and their suffering.

For the glioma model, six to eight weeks old C57BL/6JRj male mice (Janvier Labs, Le Genest-Saint-Isle, France) were used for the inoculation of the GL-261 brain tumor cell line. They were caged 3-5 animals per cage in individually ventilated cages (IVCs). The animal vivarium was a specific-pathogen-free (SPF) holding room, which was temperature- and humidity-controlled (21 ±3 °C, 50 ± 10%). All animals had ad libitum access to the same food and water throughout the entire study.

All procedures described in the present study had been previously approved by the Cantonal Veterinarian’s Office of Zurich (License 246/2015), and all efforts were made to minimize the number of animals used and their suffering.Briefly, a total of 20’000 cells in 2 μL PBS were injected at a speed of 1 μL/min intracranially into mice using a neurostar stereotaxic robot as described in Beffinger et al. 2019 (Beffinger, Schellhammer, Pantelyushin, & Vom Berg, 2019).

### GL-261luc cells

The murine GL-261 brain tumor cell line (syngenic to C57BL/6), was stably transfected with pGl3-ctrl and pGK-Puro (Promega) and selected with puromycin (Sigma-Aldrich) to generate luciferase-stable GL-261 cells. Cells were cultured at 37°C 5% CO2 in DMEM supplemented with 10% heat inactivated fetal calf serum, 1% L-glutamine (Thermo Fisher Scientific) and 1% Pen-Strep (Sigma).

### Surgical procedures

For glioma inoculation, 6-10 weeks old mice were anesthetized using a mixture of fentanyl (Helvepharm AG), midazolam (Roche Pharma AG) and medetomidin (Orion Pharma AG). GL261 cells were injected intracranially (i. c.) in the right hemisphere using a stereotactic robot (Neurostar). Briefly, a blunt-ended syringe (Hamilton; 75N, 26s/2”/2, 5 μl) was placed 1.5 mm lateral and 1 mm frontal of bregma. The needle was lowered into the burr hole to a depth of 4 mm below the dura surface and retracted 1 mm to form a small reservoir. Injection was performed in a volume of 2 μl at 1 μl/min. After leaving the needle in place for 2 min, it was retracted at 1 mm/min. The burr hole was closed with bone wax (Aesculap, Braun) and the scalp wound was sealed with tissue glue (Indermil, Henkel). Anesthesia was interrupted using a mixture of flumazenil (Labatec Pharma AG) and Buprenorphine (Indivior Schweiz AG), followed by injection of atipamezol 20 minutes later (Janssen). Carprofen (Pfizer AG) was used for perioperative analgesia. Mice were collected for experiments at 20 days post glioma inoculation, and the tumor bearing hemispheres where processed with either mechanical dissociation at 4°C or with enzymatic digestion at 37°C to test the respective technique’s efficiency in producing single cell suspensions for FACS analysis (see below).

### Brain Dissociation and Cell Isolation

The protocol has been optimized from our previous protocol (Mattei et al., 2017) and from the procedures provided by Miltenyi (adult brain dissociation kit, ABDK, Miltenyi). The protocol is optimized for the dissociation and isolation of cells from brain tissue of small volume (e.g., mouse hippocampi). A detailed dissociation protocol for larger pools of tissue or whole adult mouse brain is provided in Additional File 1. A detailed list of materials and reagents recommended for the following cell isolation protocol is displayed in Table 1 below.

**Table 1:** List of material and reagents for the cell isolation procedure.

| Material | Product Code |
| --- | --- |
| 20 ml Syringes, Bbraun | 4606205V |
| 23 G needles, 25 mm Sterican | 4657667 |
| Petri dishes, Thermo Fisher Scientific | 11339283 |
| 5 ml Tubes, Eppendorf | 0030119452 |
| 1x DPBS, Thermo Fisher Scientific | 14190144 |
| 10x DPBS, Thermo Fisher Scientific | 14200075 |
| Percoll, GE Healthcare | 17089101 |
| 1 ml Dounce Homogenizer, Active Motif | 40401 |
| Hibernate A, Brainbits | HAPR |
| 70 µm cell strainers, Fisher Scientific | 15370801 |
| Collagenase, Sigma Aldrich/Merck | 10269638001 |
| DNase I, Sigma Aldrich/Merck | 10104159001 |
| RPMI medium, Sigma Life Science | R8758-500ml |
| MACS MS Columns, Miltenyi | 130-042-201 |
| CD11b magnetic beads, Miltenyi | 130-093-634 |
| ACSA-II magnetic beads, Miltenyi | 130-097-678 |
| 7.5% BSA in PBS, Thermo Fisher Scientific | 11500496 |
| FBS, Thermo Fisher Scientific | 26140087 |
| Swing Bucket Centrifuge 5810, Eppendorf | 5810000010 |

The animals were deeply anesthetized with an overdose of Nembutal (Abbott Laboratories, North Chicago, IL, USA) and transcardially perfused with 15 ml ice-cold, calcium- and magnesium-free Dulbecco’s phosphate-buffered saline (DPBS, pH 7.3-7.4) via a 20 ml syringe and a 23 G needle (25 mm length). The brains were quickly removed and washed with ice-cold DPBS, after which the hippocampi were dissected on a cooled petri dish and placed in ice-cold Hibernate-A medium.

For enzymatic digestion (ED) at 37°C, 2-4 hippocampi/sample were grossly minced with surgical scissors and placed in a sterile 12-well plate containing dissection medium pre-heated to 37°C. The dissection medium contained Roswell Park Memorial Institute (RPMI) medium, 10% fetal bovine serum (FBS), 0.4mg/ml collagenase and 2mg/ml DNaseI. The plate was then placed in a standard cell culture incubator at 37°C with 95% O_2_ and 5% CO_2_ for a total of 30 minutes. After the first 15 minutes of incubation, the tissue was mixed with a pipette to optimize the enzymatic digestion. At the end of the 30 minutes enzymatic digestion, the plate was placed on ice, and the digested tissue was transferred to a 1 ml Dounce homogenizer to complete the dissociation of remaining tissue pieces (on ice). Thereafter, the homogenate was sieved through a 70 μm cell strainer mounted onto a 50 ml Falcon tube and transferred into 5 ml Eppendorf tubes on ice. Eppendorf tubes made of polypropylene were used because cells show less adherence to this material as opposed to polystyrene (Reichard & Asosingh, 2019).

Mechanical dissociation (MD) at 4°C was carried out on ice, while all the solutions were kept at 4°C. 2-4 hippocampi/sample were dissociated in 1.5 ml Hibernate-A medium in a 1 ml Dounce homogenizer with a loose pestle. The 1 ml Dounce homogenizer used here (see Table 1) has enough capacity for a volume of 1.5 ml, which is preferable for optimal tissue dissociation. Furthermore, this mechanical homogenizer allows enough space between the glass pot and the loose pestle, which is required for efficient tissue homogenization without extensive cell loss. The tissue was gently dounced until no bigger tissue pieces were visible. The homogenized tissue was then sieved through a 70μm cell strainer mounted onto a 50 ml Falcon tube. The Dounce homogenizer was then washed twice with 1 ml Hibernate-A, whereby each wash is poured onto the cell strainer. The homogenized hippocampi were then transferred to 5 ml Eppendorf tubes and kept on ice as described above.

After ED at 37°C or MD at 4°C, all samples were handled the same way and further processed as follows:

The homogenates were pelleted at 400xg for 6 minutes at 4°C in a swing-bucket rotor centrifuge (Eppendorf). The supernatants were removed and 1 ml ice-cold DPBS (pH 7.3-7.4) was added to all samples. The pellets were then re-suspended with a P1000 micropipette, applying a pipette-tip cut-off. After re-suspension, the final volume in each tube was brought to 1.5 ml. 500 μL of freshly prepared isotonic percoll solution was then added to each sample (final volume: 2 ml) and mixed well (applying a pipette-tip cut-off to optimize the mixing). Percoll was rendered isotonic by mixing 1 part of 10x calcium- and magnesium-free DPBS (pH 7.3-7.4) with 9-parts of percoll. Importantly, the pH of percoll was adjusted to 7.3-7.4 with 5 molar hydrochloric acid before starting the isolation procedure. The percoll solution was mixed properly with the cell suspension, after which 2 ml of DPBS were gently layered on top of it with a pipette boy set on the slowest speed, creating two separate layers. The samples were centrifuged for 10 minutes at 3000xg. The centrifugation resulted in an upper layer consisting of DPBS and a lower layer consisting of percoll. The two layers were separated by a disk of myelin and debris, while the cells were located at the bottom of the tube. The layers were aspirated, leaving about 500 μL as some cells, depending on their size, usually float in percoll just above the pellet. The cells were then washed once in DPBS making sure not to resuspend the pellet. This was achieved by gently adding 4 ml DPBS, closing the tube and holding it in a horizontal position, and gently tilting it 145 degrees in order to mix the remaining percoll with the added DPBS. The cells were then pelleted by centrifuging them at 400xg for10 minutes at 4°C.

In Figure S7 we show the utility and effectiveness of the chosen brain dissociation and cell isolation protocol in obtaining a large amount of microglia and astrocytes from as little as 1 single hippocampus from 1 brain hemisphere of adult male mice. Indeed, enough RNA can be obtained from either microglia or astrocytes from 1 hippocampus to perform scSeq, or to generate an RNA library for bulk sequencing (Figure S7).

We further compared the two protocols for their efficiency in producing microglial single cell suspensions and their proportion of live cells via flow cytometry (Figure S6b, c). We found that MD yielded a higher proportion of microglia live cells (Figure S6b) and a higher proportion of microglia singlets (Figure S6c) as compared to ED. We further compared the total amount of single cells that can be obtained from adult mouse glioma samples via our MD as compared to ED. Our MD protocol yielded a higher proportion of singlets (total cells) from tumor-bearing brain hemispheres (Figure S6d). When comparing the percentage of microglial singlets and live cells from tumor bearing hemispheres, we found that the two techniques gave comparable results (Figure S6e). Finally, as displayed in Figure 1B, when using the same number of input cells, a higher proportion of adult mouse hippocampal neurons can be obtained via this method, while a huge proportion of the latter die when using enzymatic digestion.

**Figure 1.**
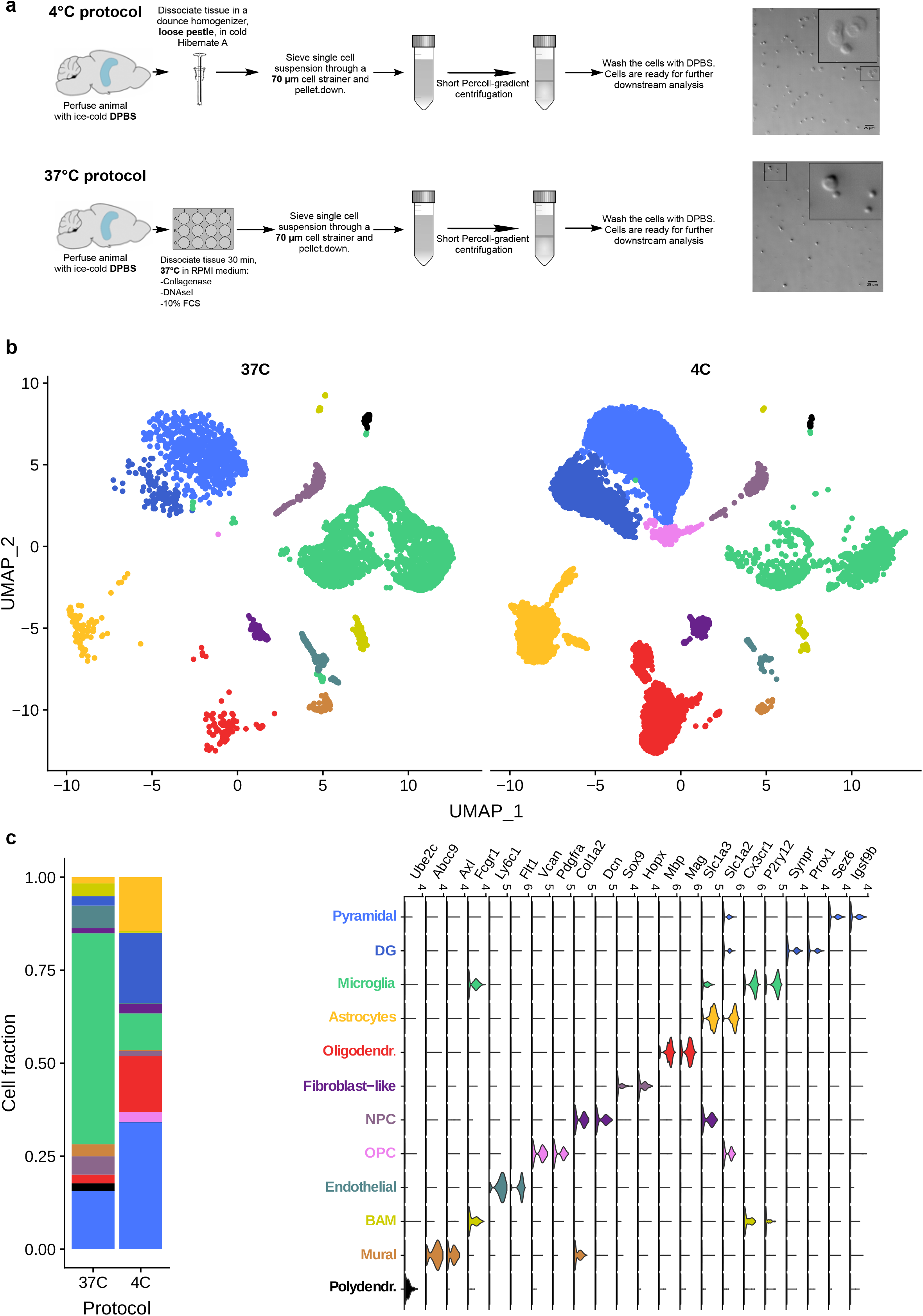
The cell isolation method affects the proportions of murine hippocampal cell populations. **(a)** Schematic illustration of the two cell isolation methods. Adult murine hippocampi were mechanically dissociated at 4°C (MD) or digested enzymatically for 30-min at 37°C (ED). After myelin removal, total hippocampal cells from both conditions were subjected to scRNA-seq. The photomicrographs show the shape of cells isolated via MD or ED. Note that cells isolated via MD appear larger and display discernible nuclei and cytoplasm. **(b)** Uniform manifold approximation and projection (UMAP) scores showing the clustering of cells isolated via ED (4448 cells) or MD (11868 cells). **(c)** Coloured legends (left) represent the identities and total fractions for each cell type and the violin plots (right) show the cell identity based on the enriched expression of specific genes.

### Microglia and Astrocyte Isolation

A schematic illustration of the procedures used for microglia and astrocyte isolation is provided in Figure 3a. For the proteomic analysis, microglia or astrocytes were isolated from 4 hippocampi (2 mice/sample) via magnetic-activated cell sorting (MACS) using mouse anti-CD11b (for microglia) or anti-ACSA-2 (for astrocytes) magnetic microbeads (Miltenyi, see Table 1) according to the manufacturer’s instructions with some modifications. The MACS buffer used consisted of 1.5% bovine serum albumin (BSA) diluted in DPBS from a commercial 7.5% cell-culture grade BSA stock (Thermo Fisher Scientific). For the isolation of astrocytes, total hippocampal cell pellets after percoll (see above) were re-suspended in 80 μL MACS buffer and 10 μL FcR-blocking reagent (Miltenyi). The cells were then incubated for 10 min at 4°C. Thereafter, 10 μL of anti-ACSA-2 microbeads were added and the cells were incubated for 15 min at 4°C. The cells were then washed with 1 ml MACS buffer and pelleted at 300xg for 5 min at 4°C. The cells were then passed through an MS MACS column attached to a magnet. This led ACSA-2-labeled cells to stay attached to the column, whereas unlabeled cells flowed through the column. After washing the columns three times with MACS buffer, astrocytes were flushed from the column with 1 ml MACS buffer and pelleted at 300xg for 5 min at 4°C. Cell pellets were then snap-frozen in liquid nitrogen and stored at −80°C. The same procedures (including the FcR-blocking step) were used to isolate microglia via anti-mouse CD11b microbeads. Figure S7b, d show a qRT-PCR-based ascertainment of microglial or astrocytic cell enrichment after MACS from total brain cells obtained with the 4°C protocol described above (See Additional File 1 for the detailed cell isolation protocol). Cells were counted manually using a standard hemocytometer (NanoEnTek, product code DHC-N01).

### RNA Extraction and Quantification

The RNA was extracted via the Lexogen Split-RNA extraction kit (Lexogen) according to the manufacturer’s instructions. The kit is based on phenol-chloroform extraction in acidic conditions, and we found this technique to yield the highest amount of RNA. RNA concentrations were measured via Qubit 4 fluorometer (Invitrogen), using the RNA HS Assay kit (Invitrogen).

### Quantitative Real-Time PCR

RNA was analyzed by TaqMan qRT-PCR instrument (CFX384 real-time system, Bio-Rad Laboratories) using the iTaq™ Universal Probes One-Step Kit for probes (Bio-Rad Laboratories). The samples were run in 384-well formats in triplicates as multiplexed reactions with a normalizing internal control. We chose 36B4 as internal standard for gene expression analyses. Thermal cycling was initiated with an incubation at 50°C for 10 min (RNA retrotranscription) and then at 95°C for 5 min (TaqMan polymerase activation). After this initial step, 39 cycles of PCR were performed. Each PCR cycle consisted of heating the samples at 95°C for 10 s to enable the melting process and then for 30 s at 60°C for the annealing and extension reaction. Cutsom-made primers with probes for TaqMan were purchased from Thermo Fisher: housekeeping gene: 36B4, product code: NM_007475.5, Siglech, prduct code: Mm_00618627_m1, P2ry12, product code: Mm00446026_m1, Gfap, product code: Mm01253033_m1, Slc1a3, product code: Mm00600697_m1. Relative target gene expression was calculated according to the Delta C(T) method.

### Single-cell RNA sequencing (scRNA-seq) using 10X Genomics platform

The quality and concentration of the single cell preparations were evaluated using an haemocytometer in a Leica DM IL LED microscope and adjusted to 1,000 cells/μl. 10,000 cells per sample were loaded into the 10X Chromium controller and library preparation was performed according to the manufacturer’s indications (single cell 3’ v3 protocol). The resulting libraries were sequenced in an Illumina NovaSeq sequencer according to 10X Genomics recommendations (paired-end reads, R1=28, i7=8, R2=91) to a depth of around 50,000 reads per cell. The quality check outcomes are displayed in Figure S2.

### Single-cell RNA-Seq Analysis

10x Chromium data was demultiplexed with cellranger mkfastq v2.0.2.Gene-cell count matrix was generated by cellranger count v2.0.2 against Ensembl GRCm38.p5 reference genome.

Single Cell RNA-Seq data analysis was carried out with R Seurat (V3) package. Specifically, to account for potential batch effects, we used canonical correlation analysis (utilized in IntegrateData function) that identifies a linear combination of features to construct a shared correlation structure and align the global transcriptome across 2 datasets (Butler, Hoffman, Smibert, Papalexi, & Satija, 2018; Stuart et al., 2019). Only genes that were detected in at least 5 cells were included in the analysis.. Furthermore, we excluded from the analysis all cells with less than 200 or more then 5000 genes detected and cells that have more than 25% mitochondrial fraction. We carried out standard preprocessing (log-normalization), and identified the top 2000 variable features for each dataset (37C and 4C). We then identified integration anchors using the FindIntegrationAnchors function. Here we used all default parameters for identifying anchors between the two data sets, setting the ‘dimensionality’ to 1:15. The final batch-corrected expression matrix was created from this anchor-set using IntegrateData function where the number of PCs used for weighting was set to 15. We employed a standard Seurat workflow for clustering and visualization: data scaling, PCA analysis and UMAP clustering using PCA reduction. For improved visualization, we further filtered out all cells located more than 3 standard deviations away from their cluster center and those which had a different identity than the majority of their 10 nearest neighbors in the UMAP.

The differential expression analysis was carried out using FindMarker function (logfc.threshold=0). P-values were calculated using Wilcoxon Rank Sum test, and adjusted with Bonferroni correction

### Code Availability

A detailed list of commands used to analyze the single cell sequencing data as well as gene count matrixes are available at https://github.com/bihealth/SC-RNA-Seq-37vs4.

### Proteotype Analysis

#### □ Liquid chromatography–tandem mass spectrometry (LC-MS/MS) analysis

The samples used for proteotype analysis were prepared using S-trap (Protifi) columns according to the manufacturer’s instructions. For MS analysis, peptides were reconstituted in 5% acetonitrile and 0.1% formic acid containing iRT peptides (Biognosys) as described in Escher et al., 2012 (Escher et al., 2012).

The peptides resulting from the isolation of microglia were analyzed in data independent acquisition (DIA) and data dependent acquisition (DDA) mode for spectral library generation. For spectral library generation, a fraction of the samples originating from the same condition were pooled to generate mixed pools for each condition. Peptides were separated by reverse-phase chromatography on a high-pressure liquid chromatography (HPLC) column (75-μm inner diameter; New Objective) packed in-house with a 50-cm stationary phase ReproSil-Pur 120A C18 1.9 μm (Dr. Maisch GmbH) and connected to an EASY-nLC 1000 instrument equipped with an autosampler (Thermo Fisher Scientific). The HPLC was coupled to a Fusion mass spectrometer equipped with a nanoelectrospray ion source (Thermo Fisher Scientific). Peptides were loaded onto the column with 100% buffer A (99% H2O, 0.1% formic acid) and eluted with increasing buffer B (99.9% acetonitrile, 0.1% formic acid) over a nonlinear gradient for 120min. The DIA method (Bruderer et al. 2017) (Bruderer et al., 2017) contained 26 DIA segments of 30,000 resolution with IT set to 60ms, AGC of 3×106, and a survey scan of 120,000 resolution with 60 ms max IT and AGC of 3×106. The mass range was set to 350-1650 m/z. The default charge state was set to 2. Loop count 1 and normalized collision energy was stepped at 27. For the DDA, a 3s cycle time method was recorded with 120,000 resolution of the MS1 scan and 20 ms max IT and AGC of 1×106. The MS2 scan was recorded with 15,000 resolution of the MS1 scan and 120 ms max IT and AGC of 5×104. The covered mass range was identical to the DIA.

The peptides resulting from the isolation of astrocytes were analyzed in DIA and DDA mode for spectral library generation. For spectral library generation, a fraction of the samples originating from the same condition were pooled to generate mixed pools for each condition. Peptides were separated by reverse-phase chromatography on a 50cm EASY-Spray C18 LC column (Thermo Fisher Scientific) connected to an EASY-nLC 1200 instrument equipped with an autosampler (Thermo Fisher Scientific). The HPLC was coupled to a Fusion Lumos mass spectrometer equipped with a nanoelectrospray ion source (Thermo Fisher Scientific). Peptides were loaded onto the column with 100% buffer A (99% H2O, 0.1% formic acid) and eluted with increasing buffer B (80% acetonitrile, 0.1% formic acid) over a nonlinear gradient for 120min. The DIA method (Muntel et al., 2019) contained 40 DIA segments of 30,000 resolution with IT set to 55ms, AGC of 1×106, and a survey scan of 120,000 resolution with 50 ms max IT and AGC of 5×105. The mass range was set to 350-1650 m/z. The default charge state was set to 2. Loop count 1 and normalized collision energy was set to 27. For the DDA, a 3s cycle time method was recorded with 120,000 resolution of the MS1 scan and 25 ms max IT and AGC of 5×105. The MS2 scan was recorded with 15,000 resolution of the MS1 scan and 35 ms max IT and AGC of 2×104. The covered mass range was identical to the DIA.

#### □ Data analysis DIA LC-MS/MS

LC-MS/MS DIA runs were analyzed with Spectronaut Pulsar X version 12 (Biognosys) (Bruderer et al., 2017) using default settings. Briefly, a spectral library was generated from pooled samples measured in DDA (details above). The collected DDA spectra were searched against UniprotKB (UniProt Swiss-prot, Mus musculus retrieved 2018) using the Sequest HT search engine within Thermo Proteome Discoverer version 2.1 (Thermo Fisher Scientific). We allowed up to two missed cleavages and semi-specific tryptic digestion. Carbamidomethylation was set as a fixed modification for cysteine, oxidation of methionine and deamidation of arginine were set as variable modifications. Monoisotopic peptide tolerance was set to 10 ppm, and fragment mass tolerance was set to 0.02 Da. The identified proteins were assessed using Percolator and filtered using the high peptide confidence setting in Protein Discoverer. Analysis results were then imported to Spectronaut Pulsar version 12 (Biognosys) for the generation of spectral libraries.

Targeted data extraction of DIA-MS acquisitions was performed with Spectronaut version 12 (Biognosys AG) with default settings using the generated spectral libraries as previously described (Bruderer et al., 2017). The proteotypicity filter “only protein group specific” was applied. Extracted features were exported from Spectronaut for statistical analysis with MSstats (version 3.8.6) using default settings (Choi et al., 2014). Briefly, features were filtered for calculation of Protein Group Quantity as defined in Spectronaut settings, common contaminants were excluded. For each protein, features were log-transformed and fitted to a mixed effect linear regression model for each sample in MSstats (Choi et al., 2014). In MSstats, the model estimated fold change and statistical significance for all compared conditions. Significantly different proteins were determined by the threshold fold-change > 2 and adjusted p-value < 0.01. Benjamini-Hochberg method was used to account for multiple testing.

### Data Accessibility

The raw data for the single-cell sequencing will be made publicly available before publication with associated access code and web link. All mass spectrometric data and acquisition information were deposited to the ProteomeXchange Consortium (www.proteomexchange.org/) via the PRIDE partner repository (Perez-Riverol et al., 2019) (data set identifier: PXD015592, username, password: kRa2alY7).

### Gene Ontology Analysis

The functional enrichment analysis for the transcriptome data was carried out with R tmod package (Zyla et al., 2019), using Utest. For each cell type we used genes that were detected by Seurat::FindMarker (logfc.threshold=0) function and sorted them by adjusted P-values: microglia – 5631 genes, astrocytes – 4108 genes, and neurons – 3288 genes. The enrichment analysis is done using GO gene set collection from MsigDB. To match MsigDB gene symbols, mouse gene ids were converted to upper case. Significantly enriched gene sets remain significant if hypergeometric test is used, for example via GOrilla, (data not shown). Significantly different genes were determined by the threshold: log-fold change > 0.5, adjusted *p*-value cut-off < 0.01. Table S1 contains a full list of deregulated genes for all cells and Table S2 contains the full GO-analysis of deregulated genes for each cell type analysed. For the proteomic GO analysis, we used hypergeometric test from the GOrilla online tool (Eden, Navon, Steinfeld, Lipson, & Yakhini, 2009). For the proteomic GO analysis, we used the hypergeometric test from GOrilla online tool: all quantified microglial or astrocytic proteins were used as background, while the significantly deregulated proteins were used as target set (all as Uniprot IDs, log-fold change > 2, adjusted *p*-value cut-off < 0.01). Significantly up- and downregulated proteins were analyzed separately using the same background set of proteins. Table S3 and Table S4 contains the full list of deregulated proteins and GO analysis per cell type respectively.

### Comparison of RNA and Protein Expression Changes

In astrocytes, we quantified 3823 proteins present in both conditions (ED and MD), which were assigned to 3623 unique genes. Median log-fold changes (LFCs) were used for the comparison of RNA and protein expression changes. For microglia, we quantified 4328 proteins that were present in both conditions (ED and MD), and these were assigned to 4158 unique genes. Pearson’s correlation was used to assess the linear association between the datasets.

### Flow Cytometric Analysis

Data analysis was performed using FlowJo 10.0.x (Treestar). Populations of interest were manually pre-gated in FlowJo software. Microglia were identified as CD45_medium_ CD11b_medium_, live, single cells. After preprocessing we combined equal number of randomly selected cells from each group and visualized data using t-Distributed Stochastic Neighbor Embedding (t-SNE). The reagents used in flow cytometry are summarized in Table 2 below.

**Table 2:** List of antibodies used for the flow cytometric analysis.

| Antigen | Clone | Color | Vendor | Product # |
| --- | --- | --- | --- | --- |
| CD11b clone M1/70 | M1/70 | BV711 | Biolegend | 101242 |
| CD11b clone M1/70 | M1/70 | APC-Cy7 | Biolegend | 101226 |
| CD45 clone 30-F11 | 30-F11 | Pacific Blue | Biolegend | 103126 |
| CD45 clone 30-F11 | 30-F11 | PerCP/Cy5.5 | Biolegend | 103131 |
| FcγRI (CD64), clone X54-5/7.1 | X54-5/7.1 | BV605 | Biolegend | 139323 |
| FcγRI (CD64), clone X54-5/7.1 | X54-5/7.1 | PE-Dazzle594 | Biolegend | 139319 |
| Cd172a (SIRPα), clone P84 | P84 | PE-Cy7 | Biolegend | 144007 |
| Cd172a (SIRPα), Cd172a (SIRPα), clone P84 | P84 | Alexa Fluor 700 | Biolegend | 144021 |
| Fixable Viability Kit | N/A | AmCyan | Biolegend | 423102 |

One hippocampus/sample (from 1 brain hemisphere) was used for the fluorescence activated cell sorting (FACS) experiments. Cells were isolated as described above. After cell isolation, the samples were washed with PBS and stained for surface markers and live/dead reagent. Following washing with PBS, the cells were fixed and permeabilized with Cytofix/Cytoperm (BD, #554715), followed by staining for intracellular markers. Flow cytometry was performed on an LSR II Fortessa (equipped with 405 nm, 488 nm, 561 nm and 640 nm laser lines; special order research product, BD) with FACS Diva Software. Before acquisition, PMT voltages were manually adjusted to reduce fluorescence spillover, and single-stain controls were acquired for compensation matrix calculation.

## Results

### Enzymatic tissue digestion induces transcriptional biases in brain cells

First, we compared single cell suspensions obtained by 30-min ED at 37°C or MD at 4°C from mouse hippocampal tissue (Figure 1a; MD-protocol was optimized in-house, see material and methods and Additional File 1 for the detailed protocol). The hippocampus was selected because it represents a discrete brain region that can be dissected in a consistent way. Visual inspection of the isolated cells showed that ED considerably affected cell morphology as compared to MD, with ED-processed cells being consistently smaller in size (Figure 1a).

We then conducted single-cell RNA-sequencing (scRNA-seq) to compare the transcriptional profile of hippocampal cell suspensions generated via both techniques. Quality check for the scRNA-seq is provided in Figure S2. Clustering using specific gene-set enrichment for cell-type identification (Hamilton, White, Rees, Wheeler, & Ascoli, 2017; Saunders et al., 2018) (Figure 1b) revealed quantitative and qualitative differences in cellular subpopulations between ED and MD. It appears evident that in tissue processed at 37°C there is a higher extent of cell death among sensitive populations, e.g. neurons and astrocytes, shifting the balance towards a higher proportion of microglia cells. Indeed, the cell population ratios seen in the MD condition are closer the known biological cellular proportions (Keller, Ero, & Markram, 2018). The differential gene expression analysis showed that ED causes significant transcriptional changes in microglia (226 deregulated genes), astrocytes (290 deregulated genes) and neurons (771 deregulated genes), (Figure 2a). We also found transcriptional deregulation in other main brain cell types, including oligodendrocytes (369 genes), endothelial cells (128 genes), border associated macrophages (134 genes), neuronal precursor cells (121 genes), fibroblast-like cells (480 genes) and mural cells (223 genes) (Figure S3a, b; a complete list of differentially expressed genes is provided in Table S1). Gene ontology (GO) analysis revealed that ED induced global deregulation in genes associated with e.g. RNA-editing, translation, metabolic functions in most cells and also immune pathways deregulation in microglia cells (Figure 2c, Figure S3; Table S2 contains the complete GO-analysis). In addition, ED led to the emergence of a distinct microglial subpopulation that was barely represented in cell suspensions obtained via MD (Figure 2b). The latter subpopulation was characterized by increased expression of immediate early genes such as *Jun, Fos* (Figure 2b), and also Egr1, Hspa8 and Jund (Figure S4) reflecting an immediate microglial response to the ED. Increased immediate early gene expression along with deregulation in genes encoding for ribosomal and mitochondrial proteins (Rpl, Rps and mt-genes) was a common feature for most brain cells exposed to ED conditions (Table S1, Figure S4). Deregulation in the latter genes has also been observed in peripheral tissues following ED (van den Brink et al., 2017), confirming what we report here. These data revealed that, alongside genes that are commonly deregulated upon enzymatic digestion, there are others which are specific to individual cell subtypes, which can be accounted for only by a tissue and cell specific analysis like the present one (Table S1 and S2, Figure S4).

**Figure 2.**
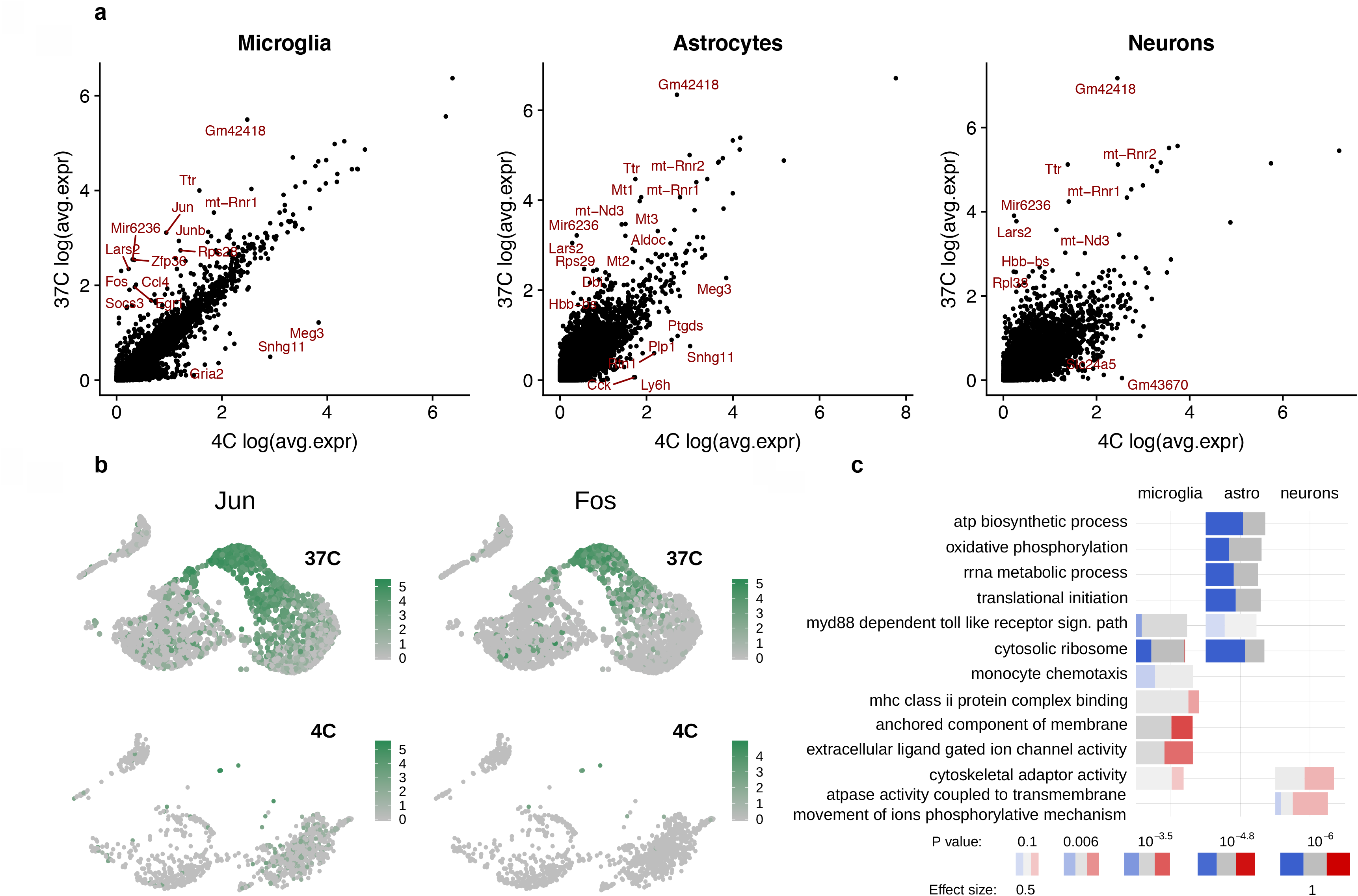
The cell isolation protocol alters the transcriptional profile of mouse hippocampal glia and neuronal cells. **(a)** Scatter plots of the differential gene expression in microglia, astrocytes, and neurons obtained via ED relative to MD. Some examples of up- and downregulated genes following ED are highlighted in red in the scatterplots. **(b)** Feature plot magnification of the microglial population from adult mouse hippocampi displaying the expression of the immediate early genes *Jun* and *Fos*. Enzymatic digestion (ED) at 37°C induces the appearance of a microglial population characterized by increased expression of immediate early genes (upper panel) which is barely present in microglia from hippocampi processed via mechanical dissociation (MD) at 4°C (lower panel). **(c)** Selected gene ontology (GO) terms associated with significantly deregulated genes in microglia, astrocytes and neurons (ED relative to MD). The bar color represents downregulation in ED relative to MD (blue) and upregulation in ED relative to MD (red). The intensity of the respective color indicates the adjusted *p*-value, while the size of the bars denotes the effect size, i.e. the area under the curve (AUC, see Fig. S3b). Significantly different genes were determined by the threshold: log-fold change > 0.5, adjusted *p*-value cut-off < 0.01.

### Enzymatic tissue digestion causes widespread proteotype artefacts in microglia and astrocytes

Despite the increasing number of studies performing proteotype analysis of brain cells freshly isolated via ED (Flowers, Bell-Temin, Jalloh, Stevens, & Bickford, 2017; Haage et al., 2019; Sharma et al., 2015), it remains unexplored whether ED introduces biological artefacts in proteotype profiles of brain cells. Therefore, we investigated whether the type of cell isolation method could also alter the proteotype of brain cells. To this end, we focused on astrocytes and microglia (Figure 3a), as they are most commonly acutely isolated for proteotype studies (Wilson & Nairn, 2018). Notably, proteotype analysis of freshly isolated brain cells is a current challenge in the field (Wilson & Nairn, 2018). We here demonstrate that our MD protocol followed by the S-trap method for protein extraction and peptide preparation enables solid mass spectrometry analyses of microglia and astrocytes extracted from only four adult mouse hippocampi (see Material and Methods).

**Figure 3.**
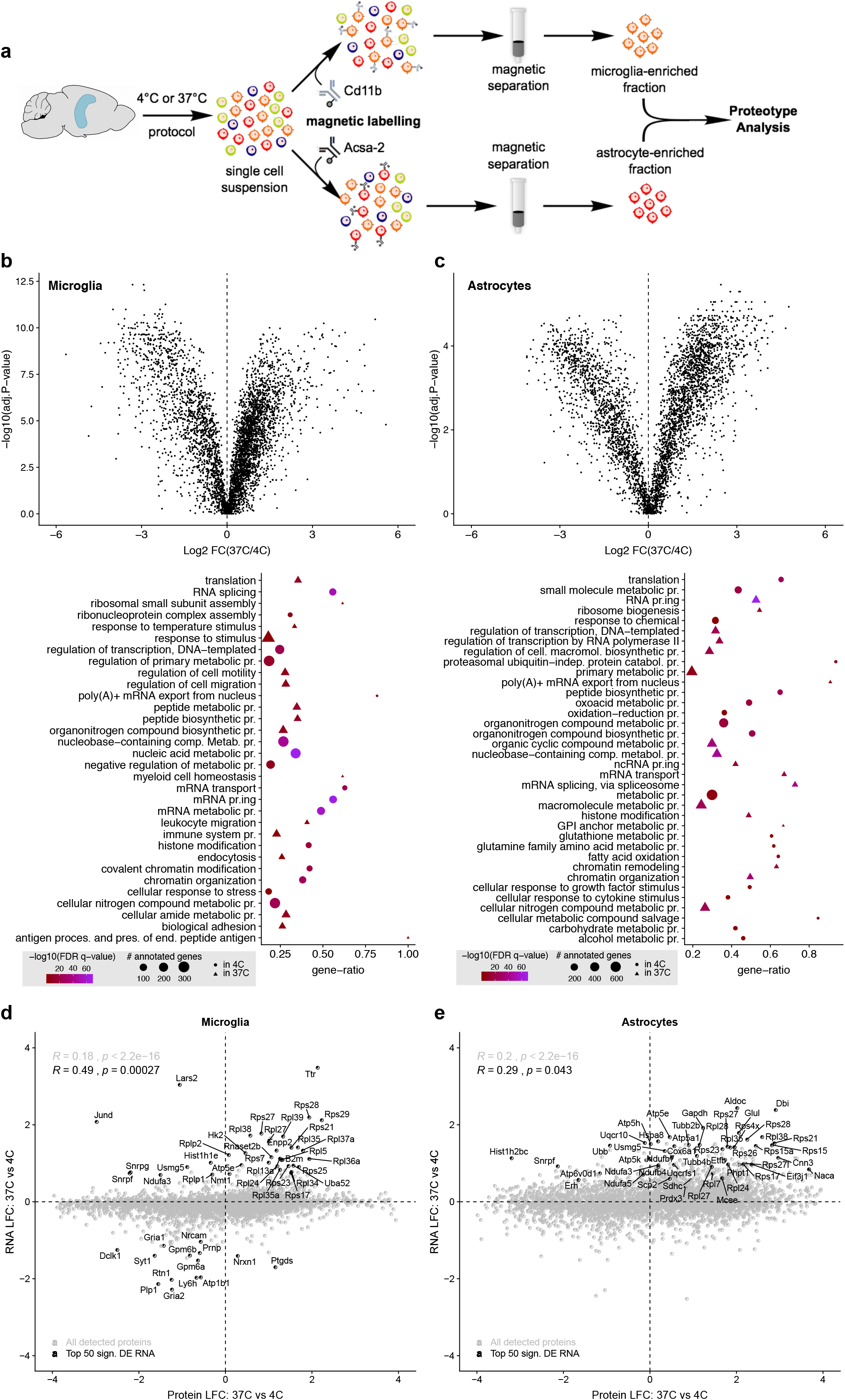
The cell isolation method affects the proteotype profiles of mouse microglia and astrocytes. **(a)** Schematic illustration of the main experimental procedure. Adult murine hippocampi were mechanically dissociated at 4°C (MD) or digested enzymatically for 30 min at 37°C (ED). After myelin removal, microglia or astrocytes where freshly isolated via magnetic associated cell sorting (MACS) for cell-specific proteotype analysis. **(b,c)** Volcano plots (top) and select gene ontology (GO) terms (bottom) of deregulated proteins in *(b)* microglia and *(c)* astrocytes obtained via ED relative to MD. N= 4 biological replicates/group. Significantly different proteins were determined by the threshold: fold-change > 2 and adjusted *p*-value < 0.01. Benjamini-Hochberg method was used to account for multiple testing. **(d,e)** Comparison of *(d)* microglial and *(e)* astrocytic RNA and protein expression differences between the two protocols. The top 50 differentially expressed genes (sorted by RNA adj. p-value), which were detected at both RNA and protein levels, are highlighted in the scatterplots. A significant correlation was detected for both microglia (R = 0.49, *p* = 0.00027) and astrocytes (R = 0.29, *p* = 0.043).

Consistent with the effects on the transcriptome (Figure 2), proteotype analysis using data-independent acquisition (DIA)-based liquid chromatography–tandem mass spectrometry (LC-MS/MS) revealed marked differences between the proteotype of microglia (Figure 3b) and astrocytes (Figure 3c) isolated via ED or MD. In microglia, 1619 proteins were significantly different after ED compared to MD. For astrocytes, we found 1984 to be significantly different following ED compared to MD. GO analysis of deregulated microglial proteins revealed that ED altered the level of proteins involved in cell motility, endocytosis, and immune processes, as well as proteins pertaining to mRNA editing, histone modifications and chromatin architecture (Figure 3b). In astrocytes, ED induced alterations in proteins associated with various metabolic processes, and with modifications in the translational and transcriptional machinery, similar to the effects on the microglial proteotype (Figure 3c).

We identified a remarkable consistency and correspondence between the effects of ED on transcriptomic and proteotype changes in both glial cell types. In fact, the top 50 deregulated RNAs significantly correlated with changes of the corresponding proteins in microglia (Figure 3d, R = 0.49, *p* = 0.00027) and astrocytes (Figure 3e, R = 0.29, *p* = 0.043). Thus, our findings show that ED is able to induce cell responses that lead to a substantial alteration in glia cell proteotype. A complete list of deregulated proteins and GO analysis is provided in Table S3 and Table S4.

Of note, in case of animal perfusion with buffers at room temperature (RT) rather than at 4°C, a thermal shift from RT to 37°C could still represent a biological shock (Mahat et al., 2016). We therefore screened whether perfusion at RT and subsequent ED at 37°C induced the same degree of proteotype alterations in microglia cells as compared to perfusion with cold buffers and subsequent ED or MD (Figure S5). We found that changes in proteotype were independent of the range of thermal shock between perfusion temperature and subsequent dissociation step at 37°C (Figure S5).

### Enzymatic tissue digestion alters the detection of classical microglial markers by flow cytometry

Finally, we also examined the influence of different cell isolation techniques on fluorescence-activated cell sorting (FACS). While ED is frequently used to produce single cell suspensions for subsequent FACS analysis of microglia (Chen et al., 2012; Martin, Boucher, Fontaine, & Delarasse, 2017; Mrdjen et al., 2018), to our knowledge no study has yet examined how cell responses during ED might influence subsequent flow cytometry analysis. Therefore, we compared whether ED, relative to MD, alters the expression of classical immune markers used in FACS-based analyses of microglia. FACS analysis confirmed that our MD protocol yielded cells of larger size than ED, as demonstrated by a significantly higher forward scatter (Figure 4a-c; gating strategy in Figure S6a). We then compared the relative expression levels of the commonly used microglial markers CD45, CD11b, SIRP-α and FcγR1 between the two methods, both on the surface and intracellularly, using differentially labelled antibodies. Surface expression was significantly increased for CD11b, CD45 and SIRP-α in ED-isolated cells compared to cells obtained via MD (Figure 4d-g). The increase in the intracellular staining of CD11b indicates a significant internalization after ED (Figure 4h, i), consistent with the increase in endocytosis-related proteins observed in the proteotype analysis. Moreover, intracellular CD11b expression was the main discriminating factor when the MD and ED conditions were clustered together (Figure 4h). Thus, the cell isolation method can influence the cellular indices used to select and study microglial cell populations in FACS-analysis. Of note, the present FACS analysis demonstrated that our MD protocol yielded a higher percentage of microglial singlets and live cells as compared ED (Figure S6b, c), showing that enzymes are not strictly necessary to obtain a successful single cell suspension.

**Figure 4.**
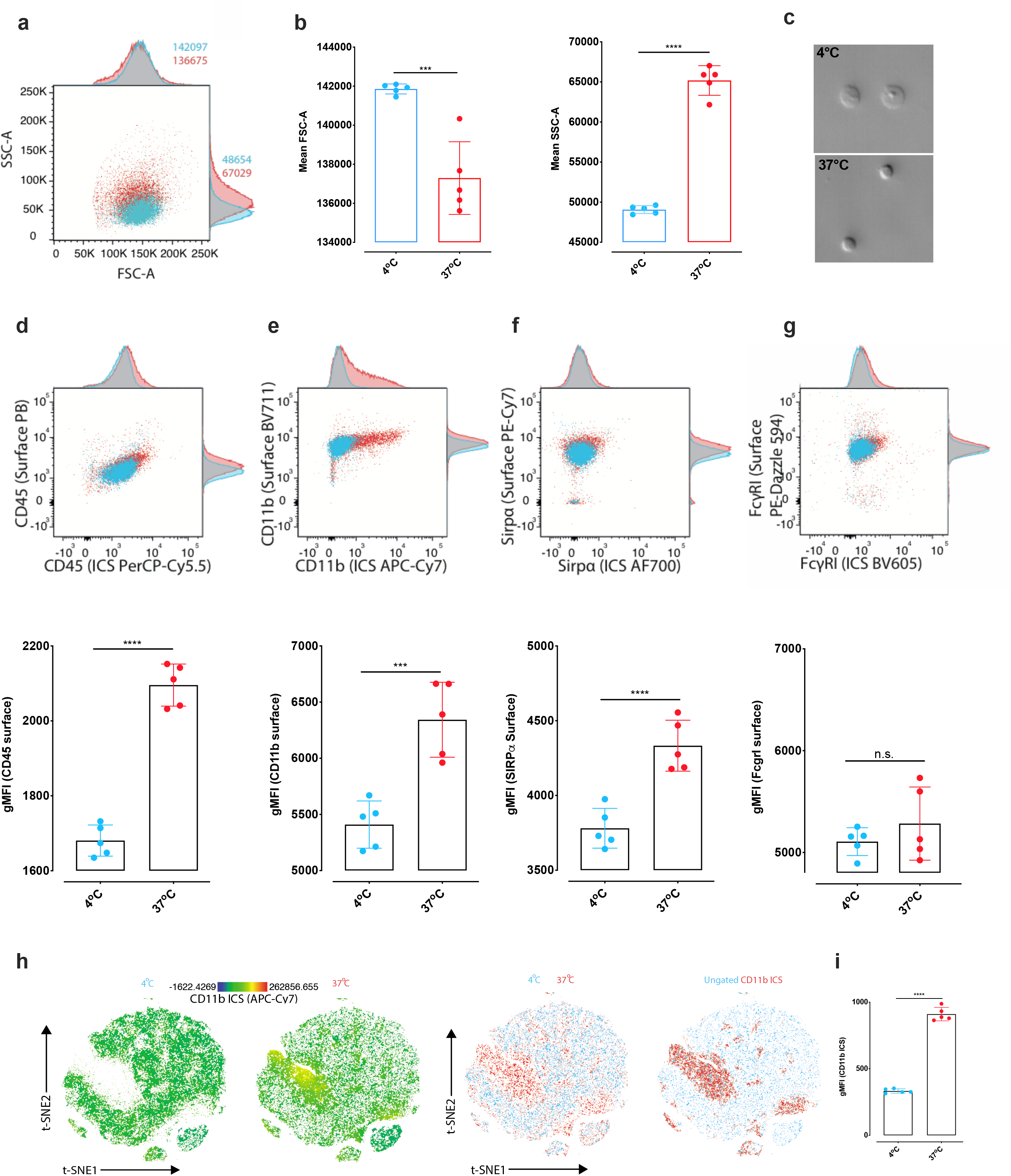
Cell isolation method affects microglial surface and intracellular marker expression in flow cytometric analysis. Flow cytometric analysis of surface and intracellular marker expression levels on microglia after enzymatic digestion (ED) at 37oC versus mechanical dissociation (MD) at 4oC. **(a)** Representative 2D-plot overlay with adjunct histograms and **(b)** scatterplots comparing forward scatter (FSC-A, t_8_ = 5.439, *p* = 0.0006) and side scatter (SSC-A, t_8_ = 18, *p* < 0.0001). **(c**) Photomicrographs illustrating the cell shape after MD or ED. **(d-g)** Representative 2D-FACS plot overlays with adjunct histograms and scatter plots comparing geometric mean fluorescence intensities (gMFIs) of surface and intracellular expression of *(D)* CD45 (t_8_ = 13.28, *p* < 0.0001), *(E)* CD11b (t_8_ = 5.288, *p* = 0.0007), *(F)* SIRPα (t_8_ = 5.213, *p* = 0.0004) and *(G)* FcγRI, (t_8_ = 1.030, *p* = 0.3). **(h)** A t-SNE clustering algorithm was used to depict CD11b expression in different populations of microglia. Left row showing relative intracellular CD11b expression levels for both conditions. Right row shows overlay of the two conditions, as well as all populations overlaid with high CD11b ICS expression. **(i)** Scatterplot for the quantification of the CD11b intracellular signal for microglia isolated via MD versus ED (t_8_ = 24.43, *p* < 0.0001) Summary of two independent experiments, N= 4 and 5 biological replicates/group respectively. Unpaired two-tailed Student *t*-tests were used to compare the means. Error bars represent the mean ± standard deviation.

## Discussion

We here provide a systematic investigation of the influence of different cell isolation protocols on transcriptional and proteotype profiles in mouse brain tissue, by taking into account single-cell transcriptomics of brain cells, proteotypes of microglia and astrocytes, and FACS analysis of microglia. Our findings indicate that ED-based cell isolation induces alterations in gene and protein expression involving both up- and downregulations. We hypothesize that such changes are due to a cellular response to the thermal shock suffered by the cells and to their response to the microenvironment present during enzymatic digestion. These alterations are likely to introduce undesirable biological biases in brain cell specific transcriptomic and proteotype profiling. While the potential risk of introducing biological artefacts through the use of ED-based cell isolation techniques has received recent attention in the context of transcriptomic studies (Ayata et al., 2018; Hrvatin et al., 2018; Wu et al., 2017), our work is the first to systematically and specifically address this issue and to extend this concern to proteotype and flow cytometry analyses. The latter is of particular importance given the increasing use of ED in mass-cytometry to study and characterize brain immune cells (Ajami et al., 2018; Bottcher et al., 2019; Mrdjen et al., 2018).

Efforts towards minimizing bias in transcriptomic datasets are indispensable not only for enhancing the reproducibility of findings, but also to avoid biological misinterpretation of datasets within studies. A way to obviate such biases is the use of single nucleus sequencing (snSeq), where whole tissue is homogenized and cells are lysed to release single nuclei that can be processed for sequencing (Bakken et al., 2018; Liu et al., 2019). This method is at present prevalently used for frozen human brain tissue samples (Habib et al., 2017), and represents a good alternative for single cell characterization of brain cells with less depth as compared to ordinary scSeq (Bakken et al., 2018). Nevertheless, snSeq is not applicable for studies requiring brain cell-specific bulk sequencing of e.g. freshly isolated microglia, to achieve a higher sequencing depth. In a few instances, scientists have adopted the use of transcription and translation blockers to avoid transcriptional and translational responses that can occur during ED (Hrvatin et al., 2018; Wu et al., 2017). However, while this strategy might be efficient in preventing *de novo* gene and protein expression, it does not prevent degradation/downregulation of pre-existing RNAs and proteins, which according to the present data, occurs during ED-based isolation as compared to MD at 4°C. Moreover, our proteotype analysis of microglia and astrocytes revealed that ED induced the deregulation of proteins associated with various epigenetic processes, including chromatin remodelling, histone modifications and DNA methylation (Figure 3, Tables S3 and S4). ED thus likely modifies the epigenetic machinery, which cannot be prevented via transcription and/or translation blockers. The latter will also not avert changes in FACS analyses of brain cells such as microglia. Indeed, here we show that ED leads to an increase in proteins associated with endocytosis (see GO table in Figure 3b) and to a marked internalization of a microglial cell marker (CD11b) as compared to MD as well as to alterations in relative expression of surface markers (Figure 4). It has to be stressed that both mechanical dissociation and ED, will alter the surface of cells due to mechanical stress and enzymatic trimming respectively, which *per se* introduces a degree of technical bias in both cases. However, holding cells in a metabolically quiescent state will prevent processes such as endocytosis and degradation that might further alter the intracellular/surface protein levels. In general, it can be argued that ED is often chosen because of difficulty in tissue dissociation, as is the case with e.g. brain tumors. We here provide evidence that the proposed mechanical dissociation protocol is efficient also with adult mouse brain tumors (Figure S6d). Human brain tumors might be more challenging and require enzymatic digestion. For these cases we would suggest enzyme mixes which work at cold temperatures such as Accutase, that can be complemented with a finishing round of Dounce homogenization as performed for the ED protocol used in the present study (see methods and Additional File 1).

Our findings corroborate the data generated by van den Brink et al. (van den Brink et al., 2017), who identified transcriptomic bias in peripheral tissue cells isolated via ED (e.g. pancreatic cells and muscle stem cells). This consistency is particularly remarkable when considering the ED-mediated deregulation in genes associated with translation, RNA processing and cellular metabolism. The latter study has been used by the authors of the *Tabula Muris* as a reference database to remove ED-induced genes bioinformatically *a posteriori* from the single cell transcriptional atlas produced in various mouse organs (Tabula Muris et al., 2018). However, the work by van den Brink et al. only considered genes that were upregulated upon ED, while not accounting for potential downregulation of genes. Our study shows that the latter can occur as well. For instance, of the 771 significantly deregulated genes in neurons, 586 were downregulated, while of the 226 significantly deregulated microglial genes, 118 were downregulated (Table S1). Likewise, according to our proteotype analysis, ED caused the downregulation of 772 proteins in microglia and of 1230 proteins in astrocytes (Table S3). Moreover, beside the general consistency between our findings and those reported by van den Brink et al., we identified several brain cell-specific alterations in biological processes after ED, which could not have been predicted by analyses of peripheral cells. A recent study by Ayata et al. also raised this issue by comparing microglia isolated via enzymatic digestion with microglial specific mRNA isolated via the translating ribosome affinity purification (TRAP) method (Ayata et al., 2018). They noted an increased expression of immediate early genes and immune related genes in their bulk sequencing of frehsly isolated microglia cells, in line with what we observe in this study. This indicates that ED-induced artifacts appear not only in scSeq, but also in bulk sequencing. In the present study, we describe an optimized protocol which provide an effective alternative to the use of transgenic mice lines involving TRAP technology or expensive dossiciation machines and enzymes (Additional File 1). The MD procedure at cold temperature here described yields a more representative picture of the cell populations due to less cell death and higher cell recovery.

In summary, although any brain cell isolation method available at present will bear a degree of technical and biological artefacts, cold MD of brain tissue helps minimize the profound alterations in the transcriptome and proteotype of specific brain cells, as seen after standard ED at 37°C. Our dataset provides a mean of identifying brain-cell specific genes and proteins affected by ED, that can be useful for correction when using previously published data sets produced via ED. Finally, we disclose a cost-effective, easily established and optimized MD protocol at 4°C which will be useful for the neuroscientific community.

## Supporting information

Additional File1_Cell_Isolation_Protocol

## Acknowledgements

This study was supported by the Swiss National Science Foundation (grant no. 310030_188524, awarded to UM; Grant No. PZ00P3_180099, awarded to JR), and by a ‘Forschungskredit’ from the University of Zurich awarded to D.M. Additional financial support was received from the European Research Council (ERC StGrant 804949 awarded to RCP) and from the Synapsis Foundation to RCP. MvO and BW were supported from ETH (grant ETH-25 15-2), the Novartis Foundation for Biomedical Research and the Swiss National Science Foundation (grant no. 31003A_160259). The authors would like to thank Dr. Emilio Yángüez and Dr. Ge Tan from the Functional Genomic Center Zurich (FGCZ) for the support in performing the single cell sequencing.

## Author contributions

D.M., A.I., R.C.P., B.W., D.B., J.v.B. and U.M. conceived and designed the study. D.M., A.I., M.v.O., S.P, J.R., F.M., M.R., M.M.B., L.S., performed the experiments and analysed the data. D.M., R.C.P., and U.M. wrote the manuscript. All authors discussed the results and commented on the manuscript.

## References

Ajami, B., Samusik, N., Wieghofer, P., Ho, P. P., Crotti, A., Bjornson, Z., … Steinman, L. (2018). Single-cell mass cytometry reveals distinct populations of brain myeloid cells in mouse neuroinflammation and neurodegeneration models. Nat Neurosci, 21(4), 541–551. doi:10.1038/s41593-018-0100-x

Ayata, P., Badimon, A., Strasburger, H. J., Duff, M. K., Montgomery, S. E., Loh, Y. E., … Schaefer, A. (2018). Epigenetic regulation of brain region-specific microglia clearance activity. Nat Neurosci, 21(8), 1049–1060. doi:10.1038/s41593-018-0192-3

Bakken, T. E., Hodge, R. D., Miller, J. A., Yao, Z., Nguyen, T. N., Aevermann, B., … Tasic, B. (2018). Single-nucleus and single-cell transcriptomes compared in matched cortical cell types. PLoS One, 13(12), e0209648. doi:10.1371/journal.pone.0209648

Beffinger, M., Schellhammer, L., Pantelyushin, S., & Vom Berg, J. (2019). Delivery of Antibodies into the Murine Brain via Convection-enhanced Delivery. J Vis Exp(149). doi:10.3791/59675

Bottcher, C., Schlickeiser, S., Sneeboer, M. A. M., Kunkel, D., Knop, A., Paza, E., … Priller, J. (2019). Human microglia regional heterogeneity and phenotypes determined by multiplexed single-cell mass cytometry. Nat Neurosci, 22(1), 78–90. doi:10.1038/s41593-018-0290-2

Bruderer, R., Bernhardt, O. M., Gandhi, T., Xuan, Y., Sondermann, J., Schmidt, M., … Reiter, L. (2017). Optimization of Experimental Parameters in Data-Independent Mass Spectrometry Significantly Increases Depth and Reproducibility of Results. Mol Cell Proteomics, 16(12), 2296–2309. doi:10.1074/mcp.RA117.000314

Butler, A., Hoffman, P., Smibert, P., Papalexi, E., & Satija, R. (2018). Integrating single-cell transcriptomic data across different conditions, technologies, and species. Nat Biotechnol, 36(5), 411–420. doi:10.1038/nbt.4096

Chen, Z., Jalabi, W., Shpargel, K. B., Farabaugh, K. T., Dutta, R., Yin, X., … Trapp, B. D. (2012). Lipopolysaccharide-induced microglial activation and neuroprotection against experimental brain injury is independent of hematogenous TLR4. J Neurosci, 32(34), 11706–11715. doi:10.1523/JNEUROSCI.0730-12.2012

Choi, M., Chang, C. Y., Clough, T., Broudy, D., Killeen, T., MacLean, B., & Vitek, O. (2014). MSstats: an R package for statistical analysis of quantitative mass spectrometry-based proteomic experiments. Bioinformatics, 30(17), 2524–2526. doi:10.1093/bioinformatics/btu305

Eden, E., Navon, R., Steinfeld, I., Lipson, D., & Yakhini, Z. (2009). GOrilla: a tool for discovery and visualization of enriched GO terms in ranked gene lists. BMC Bioinformatics, 10, 48. doi:10.1186/1471-2105-10-48

Escher, C., Reiter, L., MacLean, B., Ossola, R., Herzog, F., Chilton, J., … Rinner, O. (2012). Using iRT, a normalized retention time for more targeted measurement of peptides. Proteomics, 12(8), 1111–1121. doi:10.1002/pmic.201100463

Flowers, A., Bell-Temin, H., Jalloh, A., Stevens, S. M., Jr., & Bickford, P. C. (2017). Proteomic anaysis of aged microglia: shifts in transcription, bioenergetics, and nutrient response. J Neuroinflammation, 14(1), 96. doi:10.1186/s12974-017-0840-7

Fujita, J. (1999). Cold shock response in mammalian cells. J Mol Microbiol Biotechnol, 1(2), 243–255. Retrieved from https://www.ncbi.nlm.nih.gov/pubmed/10943555

Haage, V., Semtner, M., Vidal, R. O., Hernandez, D. P., Pong, W. W., Chen, Z., … Gutmann, D. H. (2019). Comprehensive gene expression meta-analysis identifies signature genes that distinguish microglia from peripheral monocytes/macrophages in health and glioma. Acta Neuropathol Commun, 7(1), 20. doi:10.1186/s40478-019-0665-y

Habib, N., Avraham-Davidi, I., Basu, A., Burks, T., Shekhar, K., Hofree, M., … Regev, A. (2017). Massively parallel single-nucleus RNA-seq with DroNc-seq. Nat Methods, 14(10), 955–958. doi:10.1038/nmeth.4407

Hamilton, D. J., White, C. M., Rees, C. L., Wheeler, D. W., & Ascoli, G. A. (2017). Molecular fingerprinting of principal neurons in the rodent hippocampus: A neuroinformatics approach. J Pharm Biomed Anal, 144, 269–278. doi:10.1016/j.jpba.2017.03.062

Hrvatin, S., Hochbaum, D. R., Nagy, M. A., Cicconet, M., Robertson, K., Cheadle, L., … Greenberg, M. E. (2018). Single-cell analysis of experience-dependent transcriptomic states in the mouse visual cortex. Nat Neurosci, 21(1), 120–129. doi:10.1038/s41593-017-0029-5

Keller, D., Ero, C., & Markram, H. (2018). Cell Densities in the Mouse Brain: A Systematic Review. Front Neuroanat, 12, 83. doi:10.3389/fnana.2018.00083

Liu, F., Zhang, Y., Zhang, L., Li, Z., Fang, Q., Gao, R., & Zhang, Z. (2019). Systematic comparative analysis of single-nucleotide variant detection methods from single-cell RNA sequencing data. Genome Biol, 20(1), 242. doi:10.1186/s13059-019-1863-4

Mahat, D. B., Salamanca, H. H., Duarte, F. M., Danko, C. G., & Lis, J. T. (2016). Mammalian Heat Shock Response and Mechanisms Underlying Its Genome-wide Transcriptional Regulation. Mol Cell, 62(1), 63–78. doi:10.1016/j.molcel.2016.02.025

Martin, E., Boucher, C., Fontaine, B., & Delarasse, C. (2017). Distinct inflammatory phenotypes of microglia and monocyte-derived macrophages in Alzheimer’s disease models: effects of aging and amyloid pathology. Aging Cell, 16(1), 27–38. doi:10.1111/acel.12522

Mattei, D., Ivanov, A., Ferrai, C., Jordan, P., Guneykaya, D., Buonfiglioli, A., … Wolf, S. A. (2017). Maternal immune activation results in complex microglial transcriptome signature in the adult offspring that is reversed by minocycline treatment. Transl Psychiatry, 7(5), e1120. doi:10.1038/tp.2017.80

Mrdjen, D., Pavlovic, A., Hartmann, F. J., Schreiner, B., Utz, S. G., Leung, B. P., … Becher, B. (2018). High-Dimensional Single-Cell Mapping of Central Nervous System Immune Cells Reveals Distinct Myeloid Subsets in Health, Aging, and Disease. Immunity, 48(3), 599. doi:10.1016/j.immuni.2018.02.014

Muntel, J., Kirkpatrick, J., Bruderer, R., Huang, T., Vitek, O., Ori, A., & Reiter, L. (2019). Comparison of Protein Quantification in a Complex Background by DIA and TMT Workflows with Fixed Instrument Time. J Proteome Res, 18(3), 1340–1351. doi:10.1021/acs.jproteome.8b00898

Perez-Riverol, Y., Csordas, A., Bai, J., Bernal-Llinares, M., Hewapathirana, S., Kundu, D. J., … Vizcaino, J. A. (2019). The PRIDE database and related tools and resources in 2019: improving support for quantification data. Nucleic Acids Res, 47(D1), D442–D450. doi:10.1093/nar/gky1106

Reichard, A., & Asosingh, K. (2019). Best Practices for Preparing a Single Cell Suspension from Solid Tissues for Flow Cytometry. Cytometry A, 95(2), 219–226. doi:10.1002/cyto.a.23690

Saunders, A., Macosko, E. Z., Wysoker, A., Goldman, M., Krienen, F. M., de Rivera, H., … McCarroll, S. A. (2018). Molecular Diversity and Specializations among the Cells of the Adult Mouse Brain. Cell, 174(4), 1015–1030 e1016. doi:10.1016/j.cell.2018.07.028

Sharma, K., Schmitt, S., Bergner, C. G., Tyanova, S., Kannaiyan, N., Manrique-Hoyos, N., … Simons, M. (2015). Cell type- and brain region-resolved mouse brain proteome. Nat Neurosci, 18(12), 1819–1831. doi:10.1038/nn.4160

Stuart, T., Butler, A., Hoffman, P., Hafemeister, C., Papalexi, E., Mauck, W. M., 3rd, … Satija, R. (2019). Comprehensive Integration of Single-Cell Data. Cell, 177(7), 1888–1902 e1821. doi:10.1016/j.cell.2019.05.031

Tabula Muris, C., Overall, c., Logistical, c., Organ, c., processing, Library, p., … Principal, i. (2018). Single-cell transcriptomics of 20 mouse organs creates a Tabula Muris. Nature, 562(7727), 367–372. doi:10.1038/s41586-018-0590-4

Tiklova, K., Bjorklund, A. K., Lahti, L., Fiorenzano, A., Nolbrant, S., Gillberg, L., … Perlmann, T. (2019). Single-cell RNA sequencing reveals midbrain dopamine neuron diversity emerging during mouse brain development. Nat Commun, 10(1), 581. doi: 10.1038/s41467-019-08453-1

van den Brink, S. C., Sage, F., Vertesy, A., Spanjaard, B., Peterson-Maduro, J., Baron, C. S., … van Oudenaarden, A. (2017). Single-cell sequencing reveals dissociation-induced gene expression in tissue subpopulations. Nat Methods, 14(10), 935–936. doi:10.1038/nmeth.4437

Wilson, R. S., & Nairn, A. C. (2018). Cell-Type-Specific Proteomics: A Neuroscience Perspective. Proteomes, 6(4). doi:10.3390/proteomes6040051

Wu, Y. E., Pan, L., Zuo, Y., Li, X., & Hong, W. (2017). Detecting Activated Cell Populations Using Single-Cell RNA-Seq. Neuron, 96(2), 313–329 e316. doi:10.1016/j.neuron.2017.09.026

Zhang, Y., Chen, K., Sloan, S. A., Bennett, M. L., Scholze, A. R., O’Keeffe, S., … Wu, J. Q. (2014). An RNA-sequencing transcriptome and splicing database of glia, neurons, and vascular cells of the cerebral cortex. J Neurosci, 34(36), 11929–11947. doi:10.1523/JNEUROSCI.1860-14.2014

Zyla, J., Marczyk, M., Domaszewska, T., Kaufmann, S. H. E., Polanska, J., & Weiner, J. (2019). Gene set enrichment for reproducible science: comparison of CERNO and eight other algorithms. Bioinformatics, 35(24), 5146–5154. doi:10.1093/bioinformatics/btz447

