## Supplementary Information for "Enzymatic dissociation induces transcriptional and proteotype bias in brain cell populations"

**Supporting Information**

**Supplementary Figure S1: Enzymatic digestion is the most widely used technique**

**(a)** We performed a PubMed search containing the terms "single cell sequencing, brain, 2019". Out of 172 search results we found 30 studies from 2019 that performed scSeq on fresh animal or human brain tissues. The remaining excluded studies were: unrelated search results, reviews, in vitro studies, purely bioinformatic studies and single nuclei Sequencing studies (mainly preformed on frozen human brain samples). Out of the 30 studies 23 were performed using enzymatic digestion (ED) and 7 with cold mechanical dissociation (MD). Although these are indeed not all brain scSeq studies published in 2019, this snapshot represents the general trend. **(b)** The same search for 2020 yielded 84 results of which 18 were single cell sequencing studies in animals and humans. Of these, 14 were performed via ED, 2 via MD and 2 studies did not detail the brain dissociation protocol used. The trend is similar for brain cell specific FACS and mass spectrometry analyses in published studies.

**Supplementary Figure S2: RNA-Seq QC-metrics**

**(a)** The upper panel shows the distribution of the number of genes detected per cell (left) and distribution of unique molecular identifiers (UMIs) (right). The lower panel shows mitochondrial (left) and ribosomal (right) percentage. **(b)** Variable features: Relationship between gene expression and its standard deviation. Highlighted in red are top 2000 variable genes, that are used for PCA. **(c)** Left scatter: X-axis displays the number of UMIs per cell and the Y-axis is the mitochondrial percentage per cell. Right scatter plot shows the relationship between the number of detected genes and UMIs per cell.

**Supplementary Figure S3: The cell isolation method affects the transcriptomic profiles of several hippocampal cell types in the mouse.**

Alterations in gene expression were observed in most other hippocampal cell types after 37°C enzymatic digestion (ED) relative to cold mechanical dissociation (MD). Displayed in **(a)** are select gene ontology terms associated with genes deregulated in response to ED in oligodendrocytes, neuronal precursor cells (NPC), border associated macrophages (BAM), endothelial cells, mural cells and fibroblast-like cells. The bar color represents downregulation in ED relative to MD (blue) and upregulation in ED relative to MD (red). The intensity of the respective color indicates the adjusted *p*-value, while the size of the bars denotes the effect size, i.e. the area under the curve. Depicted in **(b)** are the evidence plots showing the area under the curve for selected gene ontology terms and the genes within the latter. X axis is the gene list reported by Seurat::FindMarker function sorted by adjusted p-value. (at 0 is the gene with the lowest p-value). Y axis is the cumulative fraction of genes in a specific GO term. Higher accumulation of those genes in the top of the list (closer to 0 on X axis) results in larger AUC. The color of the curve represents the specific cell type as depicted in Fig 1.

The full list significantly deregulated genes in each cell type and the associated gene ontology terms can be found in Supplementary Tables 1 and 2 respectively.

**Supplementary Figure S4: Cell isolation method affects the expression of immediate early genes and genes associated with RNA and cellular metabolism.**

Violin plots showing the expression distribution of selected examples of significantly differentially expressed genes in different cell populations from ED and MD dissociation conditions. Consistent with what reported by van den Brink et al in peripheral tissue subjected to ED, (van den Brink et al., 2017) we find a global upregulation of the immediate early genes (*Jun, Egr1, Jund, Junb*) and heat-shock protein genes (*Hspa1a, Hspa1b, Hspa8*). We also observed a global induction of genes associated with RNA-metabolic processes such as *Snrpg* which is associated with alternative splicing functions (Papasaikas, Tejedor, Vigevani, & Valcarcel, 2015). Furthermore, represented in this figure are some examples of ED-induced downregulation of select genes. For instance, worth reporting is the downregulation of long noncoding RNAs such as *Meg3* and *Malat1*. The latter are receiving increasing attention as regulators of brain development and players in various brain diseases (Sanli et al., 2018; Wang et al., 2018; Zhang, Hamblin, & Yin, 2017). Also, we found a rather global downregulation in the proteolipid protein gene *Plp1* which holds important roles in myelin sheet stability and microglial immune responses, (Tanaka et al., 2009; Tatar et al., 2010). Remarkable downregulation following ED was observed also for *Ly6h*, which has been shown to modulate the hippocampal α7-nicotinic acetylcholine receptor subunit and consequently glutamatergic signaling (Puddifoot, Wu, Sung, & Joiner, 2015). The full list significantly deregulated genes in each cell type and the associated gene ontology terms can be found in Supplementary Tables 1 and 2 respectively.

**Supplementary Figure S5: Enzymatic digestion affects the proteotype profile of hippocampal microglia regardless of perfusion temperature.** Some research groups perfuse animals at room temperature (RT) rather than with ice-cold buffers. We sought to identify whether perfusion at RT and subsequent ED at 37°C for 30 min also leads to a significant cellular response in microglia. **(a)** The volcano plot shows the deregulated proteins in microglia cells extracted from mice perfused at RT followed by enzymatic digestion at 37°C as compared to microglial proteins from mice perfused at 4°C followed by mechanical dissociation at 4°C. We found overall 2130 proteins with significant abundance difference. The GO terms for biological processes associated with this deregulation are similar to the ones observed for the experiment in Figure 2 and they are listed in Table S3. **(b)** Volcano plot displaying the deregulated proteins in microglia cells extracted from mice perfused at RT followed by enzymatic digestion at 37°C as compared to microglial proteins from mice perfused at 4°C followed by enzymatic digestion at 37°C. Only few proteins (66) were identified as significantly different in abundance. **(c)** Heat-map showing the log2 protein abundance of microglia from mice perfused and isolated at 4°C in comparison to the profile of microglia from mice perfused at either RT or at 4°C and subsequently isolated via enzymatic digestion at 37°C. Overall, this analysis demonstrates that enzymatic digestion at 37°C causes a substantial proteomic deregulation in microglia cells, regardless of whether the animals are perfused at RT or at 4°C. Significantly different proteins were determined by the threshold fold-change > 2 and adjusted p-value < 0.01. Benjamini-Hochberg method was used to account for multiple testing. N= 4 biological replicates/group. A complete list of differentially regulated proteins can be found in Supplementary Table S3. Supplementary Table S4 contains the complete list of GO-terms associated with deregulated proteins in all conditions described.

**Supplementary Figure S6: Back gating**

**(a)** Gating strategy used for the FACS analysis of microglia cells. Single cells were gated followed by a gating for live-cells and gating out debris. Microglia cells were selected based on their $CD11b_{+}$ and $CD45_{low}$ expression. **(b, c)** We found that mechanical dissociation yielded a higher proportion of live microglia *(b)* ($t_8 = 18.11$, $p < 0.0001$) and microglia singlets *(c)* ($t_8 = 12.14$, $p < 0.0001$) as compared to enzymatic digestion at 37°C. *(d)* Mechanical dissociation also yielded a higher percentage of total single cells from adult mouse tumor samples ($t_7 = 4.38$, $p = 0.0032$). Unpaired two-tailed Student t test was used to compare the means. Error bars represent the mean ± standard deviation. Summary of two independent experiments, N= 4-5 biological replicates/group.

**Supplementary Figure S7: Glia cells and RNA yields following cold mechanical dissociation**.

**(a)** Table showing examples of the number of microglia and the respective total RNA that can be obtained from one single adult mouse hippocampus (from one hemisphere) or two hippocampi (from one mouse) using the mechanical dissociation protocol at 4°C proposed in this study (see methods). Microglia cells were sorted via Magnetic Associated Cell Sorting (MACS) using anti-CD11b microbeads. **(b)** The graphs show the successful enrichment of the MACS sorted microglia cells as compared to the flow through via qRT-PCR for the microglial specific genes *Siglech* and *P2ry12*. **(c)** Table showing examples of the number of astrocytes and the respective total RNA that can be obtained from one single adult mouse hippocampus (from one hemisphere) or two hippocampi (from one mouse) using the mechanical dissociation protocol at 4°C proposed in this study. Microglia cells were sorted via MACS using anti-ACSA2 microbeads. **(d)** The graphs show the successful enrichment of the MACS sorted astrocytes as compared to the flow through via qRT-

PCR for the astrocytic genes *Gfap* and *Slc1a3*. Abbreviations: ft: flow through, tc: target cells. N= 5 biological replicates/group. Error bars represent the mean ± standard deviation.

**Supplementary Table S1:** Differential gene expression for neurons, microglia, astrocytes, oligodendrocytes, endothelial cells, neuronal precursor cells, border associated macrophages, endothelial cells, mural cells and fibroblast-like cells. The differential expression analysis presented is based on a Log-fold change cut-off > 0.5 and adjusted p-value <0.01.

**Supplementary Table S2:** Complete list of gene ontology terms for biological process, function and component for the genes deregulated upon enzymatic digestion relative to cold mechanical dissociation in all analysed cell types.

**Supplementary Table S3:** Complete list of differentially regulated proteins for microglia and astrocytes isolated from enzymatically digested and mechanically dissociated hippocampal tissues. Significantly different proteins were determined by the threshold: fold-change > 2 and adjusted *p*-value < 0.01. Benjamini-Hochberg method was used to account for multiple testing.

**Supplementary Table S4:** Complete list of gene ontology terms for biological process, function and component for the proteins deregulated upon enzymatic digestion in microglia and astrocytes.

**Additional File 1:** Complete description of the 4°C brain cell isolation procedure.

#### References


Papasaikas, P., Tejedor, J. R., Vigevani, L., & Valcarcel, J. (2015). Functional splicing network reveals extensive regulatory potential of the core spliceosomal machinery. *Mol Cell, 57*(1), 7-22. doi:10.1016/j.molcel.2014.10.030

Puddifoot, C. A., Wu, M., Sung, R. J., & Joiner, W. J. (2015). Ly6h regulates trafficking of alpha7 nicotinic acetylcholine receptors and nicotine-induced potentiation of glutamatergic signaling. *J Neurosci, 35*(8), 3420-3430. doi:10.1523/JNEUROSCI.3630-14.2015

Sanli, I., Lalevee, S., Cammisa, M., Perrin, A., Rage, F., Lleres, D., . . . Feil, R. (2018). Meg3 Non-coding RNA Expression Controls Imprinting by Preventing Transcriptional Upregulation in cis. *Cell Rep, 23*(2), 337-348. doi:10.1016/j.celrep.2018.03.044

Tanaka, H., Ma, J., Tanaka, K. F., Takao, K., Komada, M., Tanda, K., . . . Ikenaka, K. (2009). Mice with altered myelin proteolipid protein gene expression display cognitive deficits accompanied by abnormal neuron-glia interactions and decreased conduction velocities. *J Neurosci, 29*(26), 8363-8371. doi:10.1523/JNEUROSCI.3216-08.2009

Tatar, C. L., Appikatla, S., Bessert, D. A., Paintlia, A. S., Singh, I., & Skoff, R. P. (2010). Increased Plp1 gene expression leads to massive microglial cell activation and inflammation throughout the brain. *ASN Neuro, 2*(4), e00043. doi:10.1042/AN20100016

van den Brink, S. C., Sage, F., Vertesy, A., Spanjaard, B., Peterson-Maduro, J., Baron, C. S., . . . van Oudenaarden, A. (2017). Single-cell sequencing reveals dissociation-induced gene expression in tissue subpopulations. *Nat Methods, 14*(10), 935-936. doi:10.1038/nmeth.4437

Wang, D. Q., Fu, P., Yao, C., Zhu, L. S., Hou, T. Y., Chen, J. G., . . . Zhu, L. Q. (2018). Long Non-coding RNAs, Novel Culprits, or Bodyguards in Neurodegenerative Diseases. *Mol Ther Nucleic Acids, 10*, 269-276. doi:10.1016/j.omtn.2017.12.011

Zhang, X., Hamblin, M. H., & Yin, K. J. (2017). The long noncoding RNA Malat1: Its physiological and pathophysiological functions. *RNA Biol, 14*(12), 1705-1714. doi:10.1080/15476286.2017.1358347

**Supplementary Figure S1**

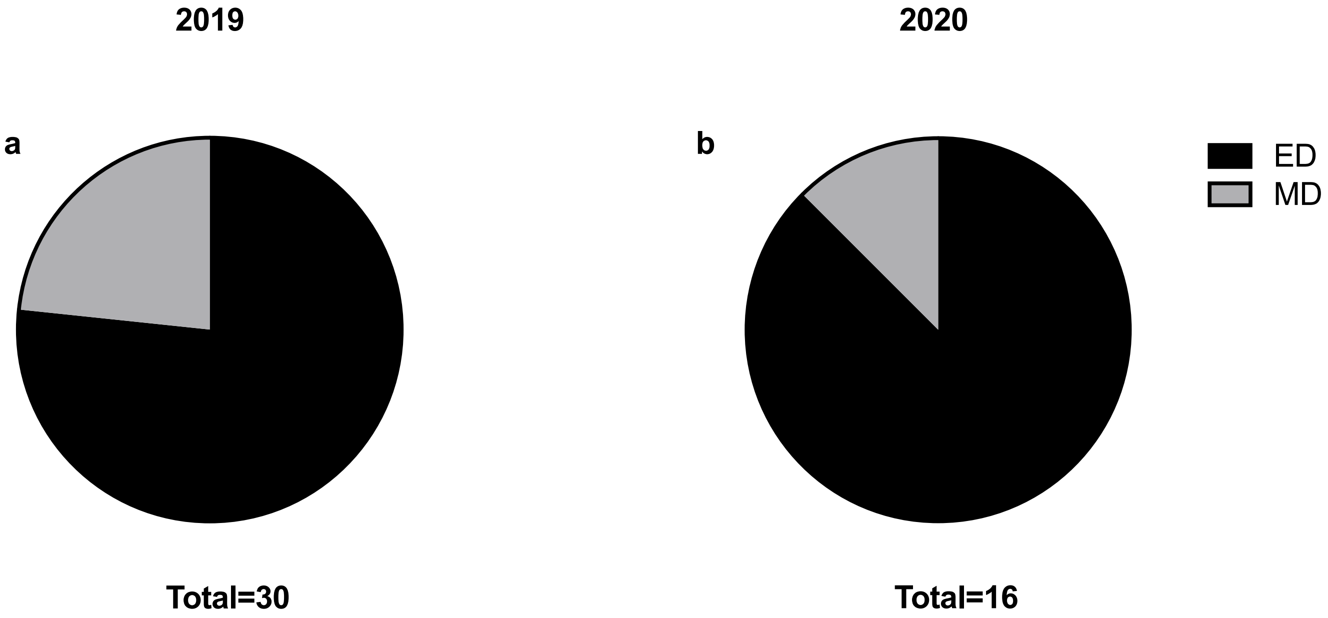

### Supplementary Figure S2

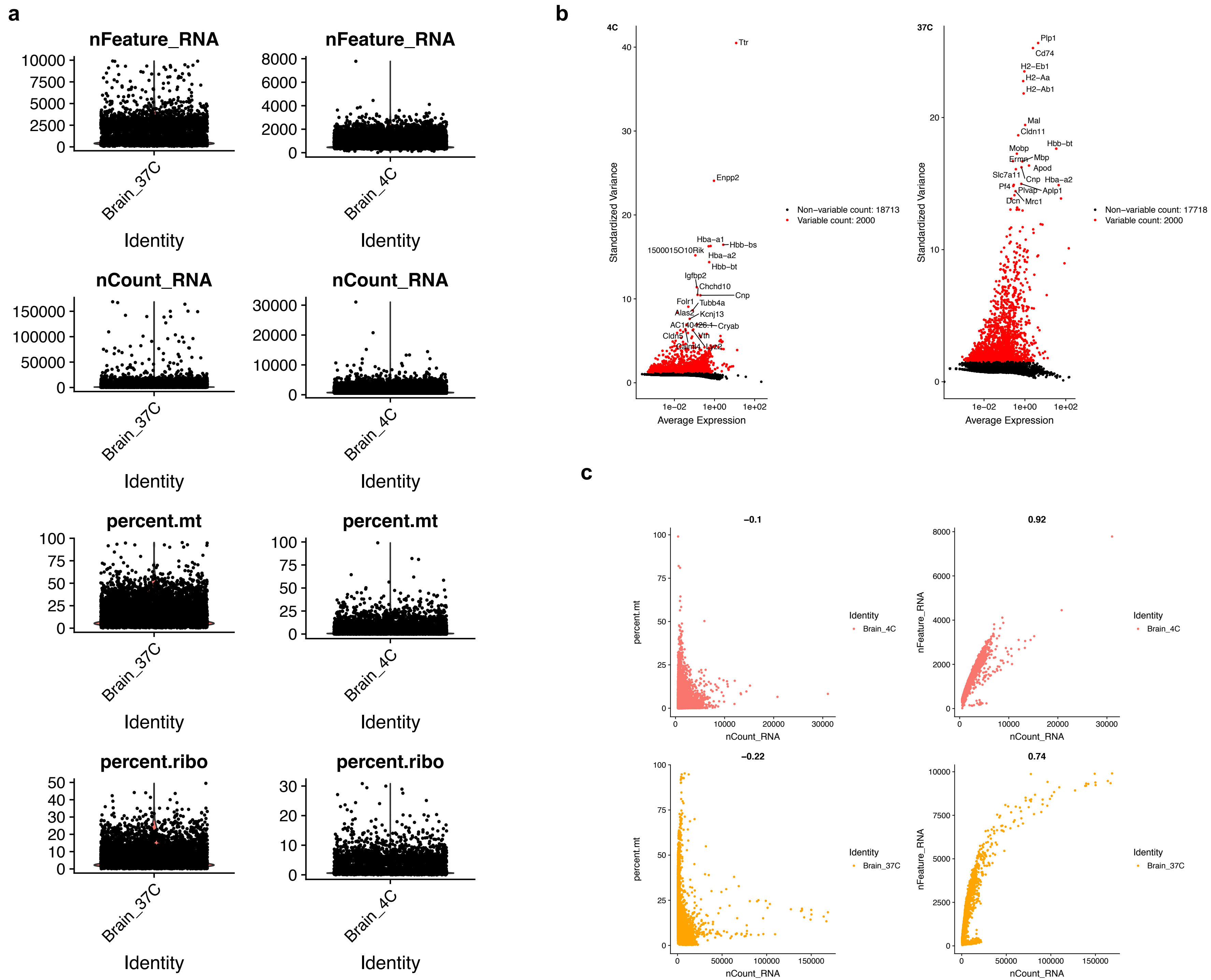

### Supplementary Figure S3

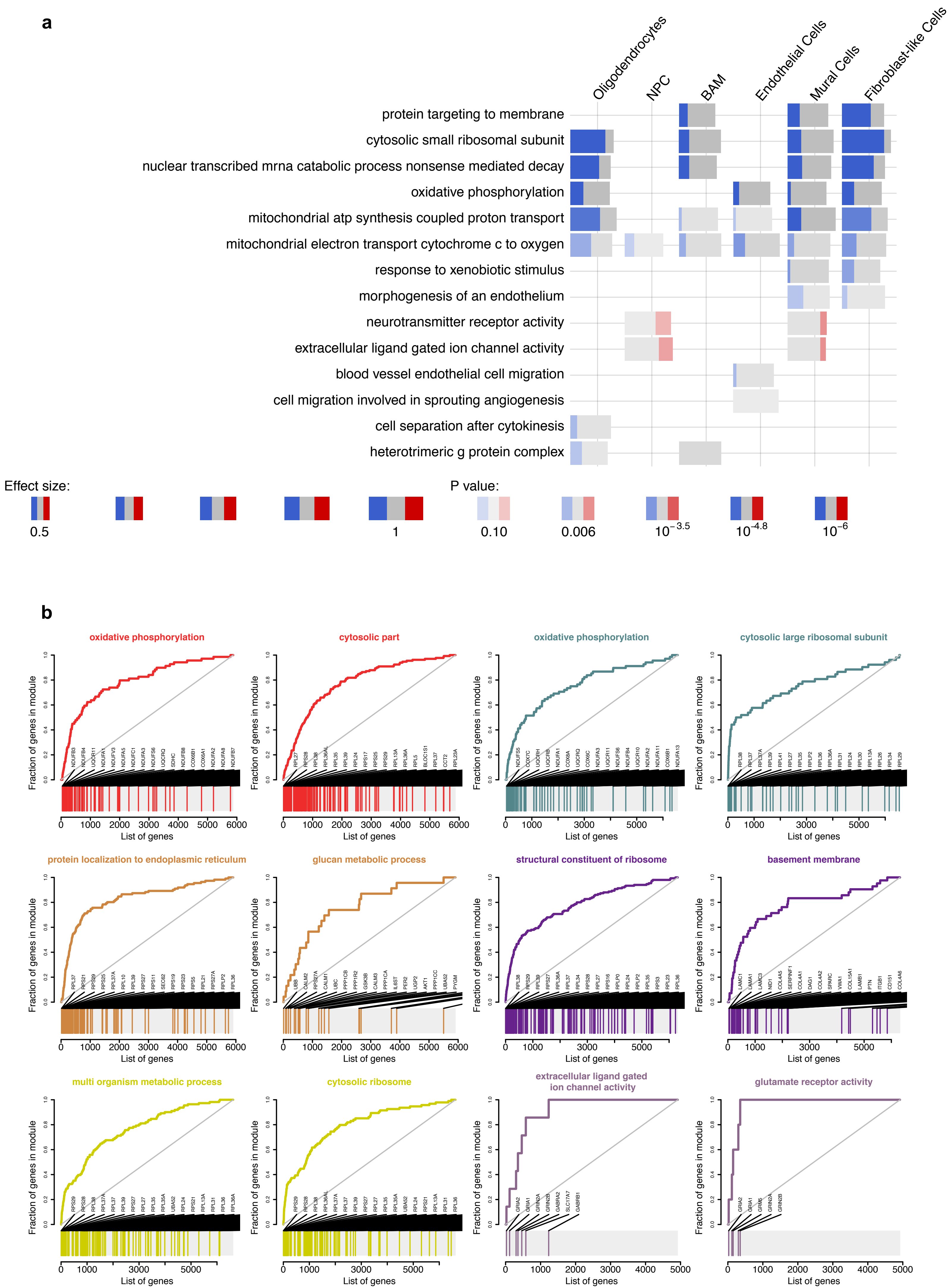

#### Supplementary Figure S4

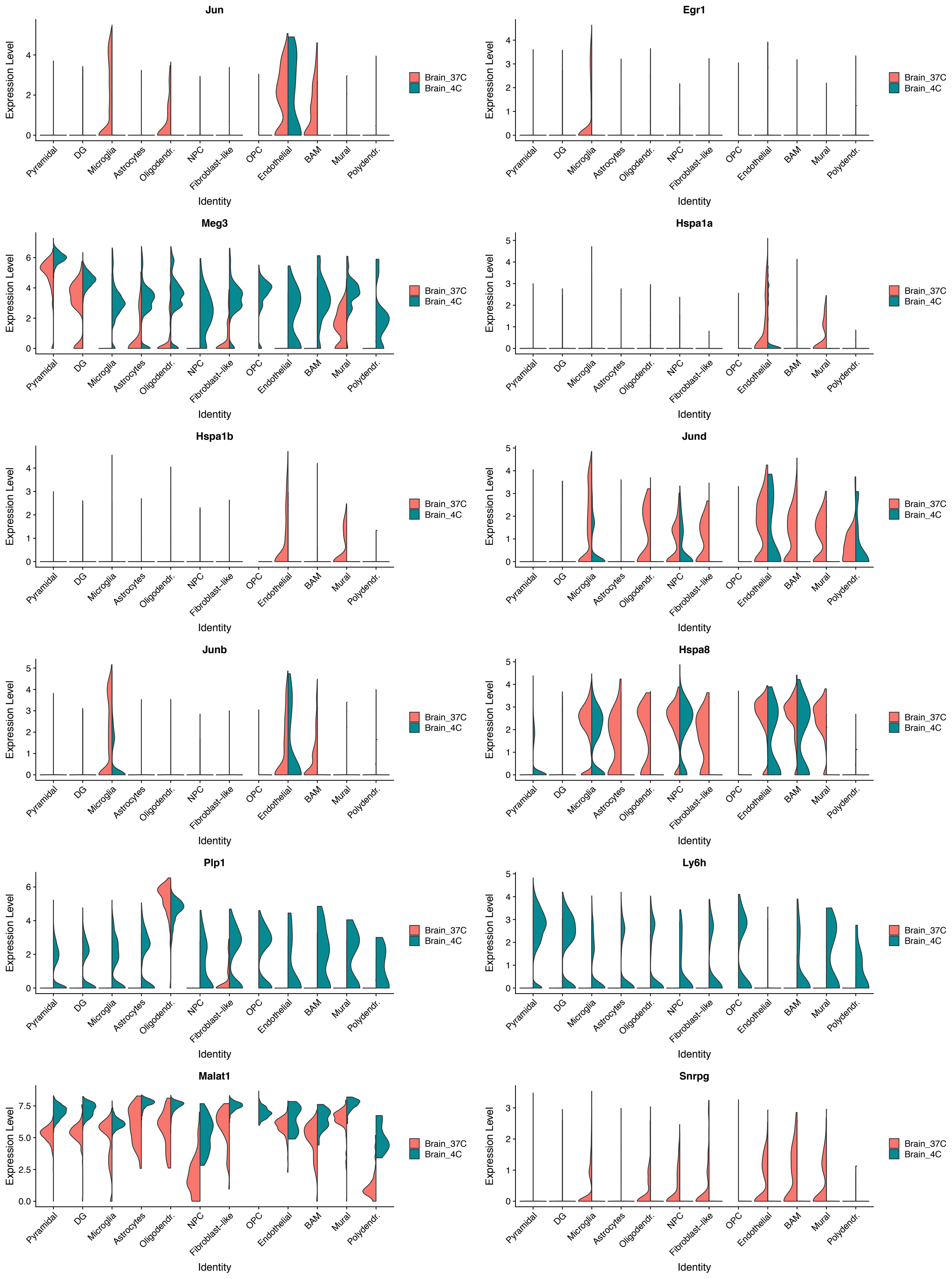

**Supplementary Figure S5**

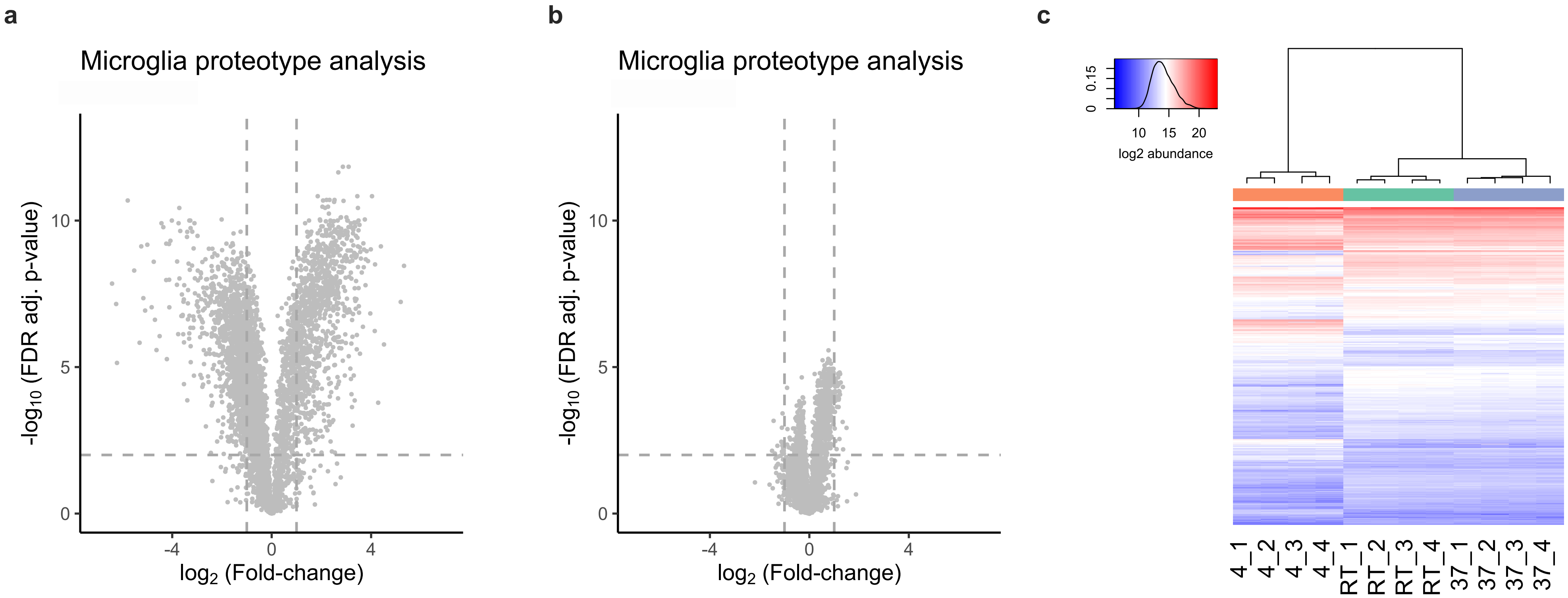

**Supplementary Figure S6**

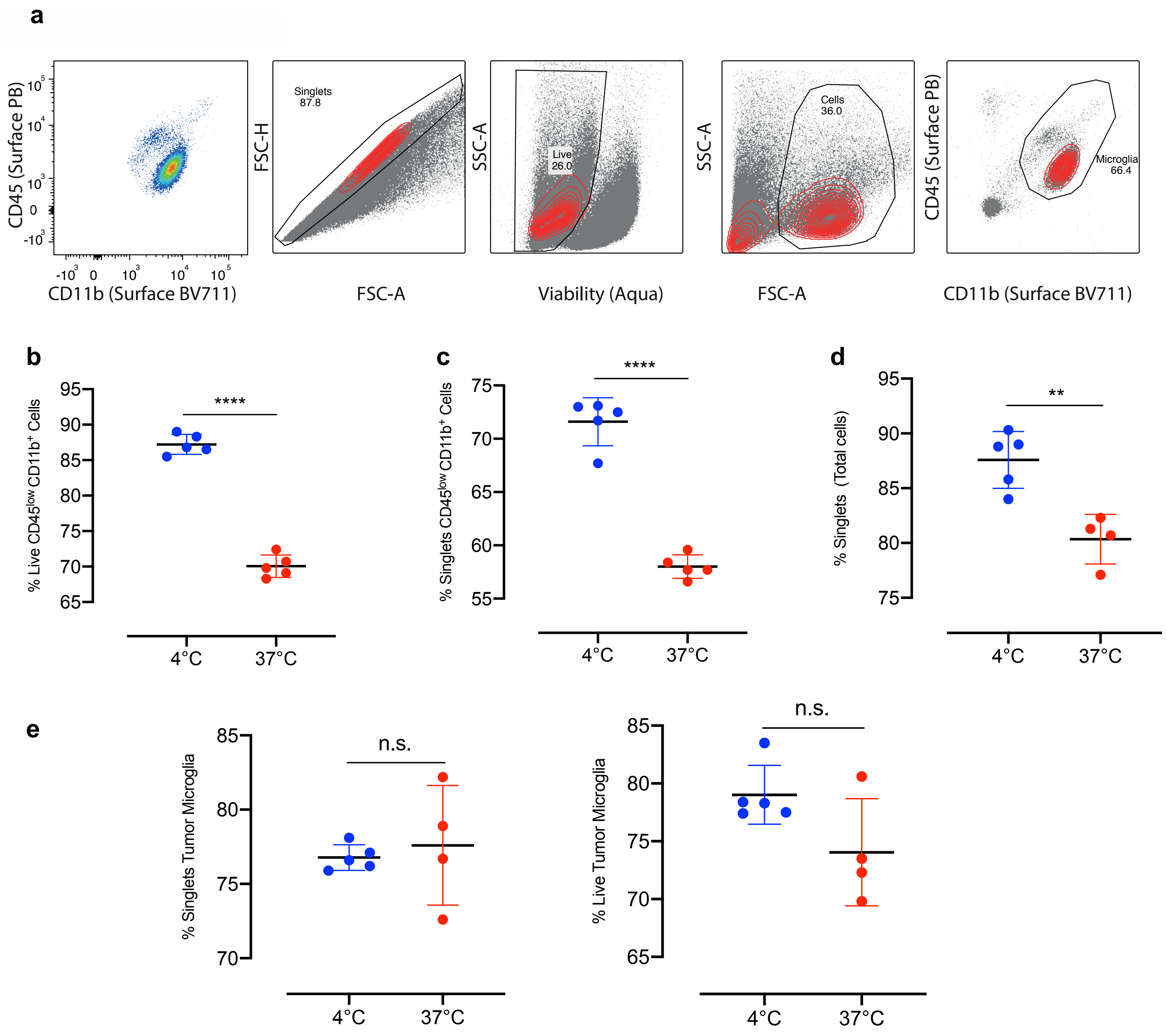

**Supplementary Figure S7**

**a**

| 1 Hippocampus | | |
|---|---|---|
| Sample | Number of Microglia | Total RNA extracted (ng) |
| 1 | 150.000 | 85 |
| 2 | 187.500 | 74 |
| 3 | 167.500 | 88 |
| 4 | 100.000 | 64 |
| 5 | 95.000 | 59 |

| 2 Hippocampi | | |
|---|---|---|
| Sample | Number of Microglia | Total RNA extracted (ng) |
| 1 | 432.500 | 214 |
| 2 | 362.500 | 197 |
| 3 | 396.000 | 203 |
| 4 | 450.000 | 218 |
| 5 | 442.500 | 231 |

**b**

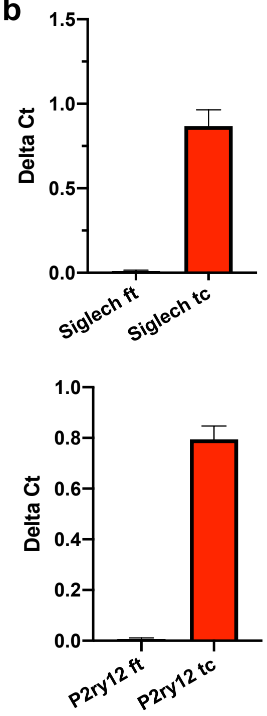


**c**

| 1 Hippocampus | | |
|---|---|---|
| Sample | Number of Astrocytes | Total RNA extracted (ng) |
| 1 | 127.500 | 104 |
| 2 | 122.500 | 99 |
| 3 | 150.000 | 108 |
| 4 | 110.000 | 80 |
| 5 | 120.000 | 97 |

| 2 Hippocampi | | |
|---|---|---|
| Sample | Number of Astrocytes | Total RNA extracted (ng) |
| 1 | 317.500 | 310 |
| 2 | 300.000 | 272 |
| 3 | 330.000 | 378 |
| 4 | 285.000 | 210 |
| 5 | 307.500 | 225 |

**d**

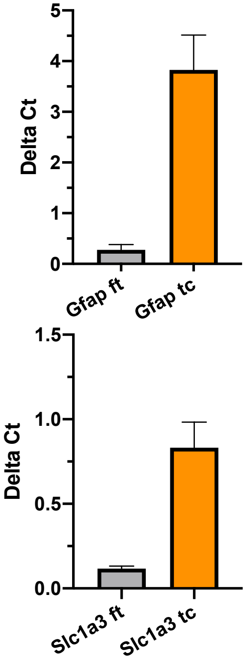
