## Additional File1_Cell_Isolation_Protocol for "Enzymatic dissociation induces transcriptional and proteotype bias in brain cell populations"

### Brain Cell Isolation Protocol

Mattei *et al.*

**To prepare before starting the procedure:**

⇒ Adjust the pH of 1x DPBS to 7.3-7.4 (stored at 4°C).
⇒ Adjust the pH of percoll to 7.3-7.4 (stored at 4°C). *See note in the last page.
⇒ Make sure the pH of 10x DPBS is 7.3-7.4 (stored at room temperature).
⇒ Prepare 5 ml Eppendorf tubes with 2 ml Hibernate-A medium on ice to collect and store the tissue.
⇒ Depending on the amount of tissue to be processed, cell isolation can be carried out in either 15 ml Falcons or in 5ml Eppendorf tubes (see Table 1).
⇒ **Use a swing bucket centrifuge set at 4°C for all centrifugation steps.** *Of Note: some swing-bucket centrifuges tend to cool further down than 4°C during centrifugation. If this is the case for your centrifuge, set the centrifuge to 6°C in order to avoid temperatures below 4°C. Excessive cold temperature can cause crystal-build-up inside the cells which will destroy them and lead to lower cell yield.*

**Procedure**

All the following steps are carried out on ice. All centrifugation steps are carried out at 4°C

**1.** Anesthetize the animal.
**2.** Perfuse intracardially with 15 mL **cold** DPBS via a 20ml syringe with a 23G needle (length: 25 mm).
**3.** Quickly remove the brain and place it on a petri-dish on ice and dissect the region(s) of interest. Place the brain region into a 5 ml tube with Hiberenate-A on ice.
**4.** Dissociate the brain region in 1.5 ml Hibernate-A in a 1 ml Dounce homogenizer using the **Loose** pestle, which shreds the tissue but preserve the cells. The number of strikes depend on the size and consistency of the specific areas, simply make sure there are no bigger tissue pieces left. *Note: Most Dounce homogenizers have the capacity for excess volumes. In this case, the 1 ml Dounce homogenizer can be filled up to 2 ml.*

**5.** *If processing half or a whole adult mouse brain: pre-mince the brain with surgical scissors and transfer the brain into the Dounce homogenizer. Homogenize in 2 ml Hibernate-A.*

**6.** Place a 70 μm strainer on a 50 ml falcon tube. **Pre-wet the strainer with 1 ml Hibernate-A** medium.
**7.** Once the tissue is shredded, pass the suspension through the strainer to collect it into the 50 ml falcon tube.
**8.** If there are tissue pieces left on the strainer that were not properly dissociated in the Dounce homogenizer: use the rubber-tip of a 1 ml-syringe to delicately dissociate it through the strainer before proceeding with the washes in step 9.

**Table 1**

| Size of Brain Region | Volume of DPBS | Volume of Isotonic Percoll | Top layer of DPBS |
|---|---|---|---|
| **Bigger tissue pieces (e.g. half-whole brain)**<br><br>**15 ml Falcon** | Add 2 ml DPBS, resuspend (cut off pipette-tip). | Add 1000 μl isotonic percoll. **check that the final volume is 4ml! If it is not, add DPBS up to 4ml.** Mix well to make the solution homogeneous (with pipette tip cut-off). | **Carefully** apply a layer of 4 ml DPBS on top with a pipette-boy set on the slowest speed.<br><br>**Final volume: 8 ml** |
| **Smaller brain areas (e.g. 2-6 Hippocampi)**<br><br>**5 ml Eppendorf tubes** | Add 1 ml DPBS, resuspend (cut off pipette-tip). | Add 500 μl isotonic percoll. **check that the final volume is 2 ml! If it is not, add DPBS up to 2 ml.** Mix well to make the solution homogeneous (with pipette tip cut-off). | **Carefully** apply a layer of 2 ml DPBS on top with a pipette-boy set on the slowest speed.<br><br>**Final Volume: 4 ml** |

9. Quickly rinse the Dounce homogenizer 2-3 times (depending on the amount of tissue used) with 1ml Hibernate-A/wash and transfer through the 70 μm strainer to collect cells that stayed on top of the strainer.

10. Using a pipette-boy, transfer the single cell suspension collected in the 50 ml falcon into pre-cooled 15 ml falcons or 5 ml Eppendorf tubes (depending on the amount of tissue being processed).

11. Centrifuge the single cell suspension at 350 rcf, 10 min (if processing bigger brain pieces using 15 ml tubes, centrifuge at 400 rcf, 10 min).
12. During this time, prepare the Isotonic Percoll: E.g. Take 9 ml pH-adjusted percoll and add 1ml 10x DPBS (pH-adjuasted), mix well. For smaller sample numbers: mix 4.5 ml Percoll with 500 μl 10x DPBS.
13. Aspirate the supernatant and resuspend the pellet in appropriate volume of cold DPBS. Resuspend the pellet with a P1000 applying a pipette-tip cut-off to ensure proper resuspension, as the pellet is thick – **See the Table 1 above and follow the indications.**
14. Centrifuge the layered samples at 3000 rcf, 10min.
15. After centrifugation: Aspirate the supernatant including the myelin disk in-between the two phases, leaving about 500μl left in the tube as there are some cells visibly floating above the pellet.
16. **Gently** add 4ml cold DPBS. **OBSERVE:** place the tip of the pipette-boy against the wall of the tube and carefully apply the DPBS **(Do not directly resuspend the pellet!)**.
17. In order to properly mix the remaining percoll and the added DPBS: gently place the tube horizontally at 180° and then carefully tilt it three times as displayed in **Figure 1** below.
18. Pellet the cells by centrifuging at 400 rcf, 10min - **This is your final pellet** (total brain cells)

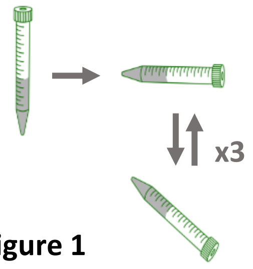


**Figure 1**

**Note:** In case of further magnetic cell sorting or flow cytometry, which require incubation steps with subsequent washes, we found that the cells can be successfully pelleted at 300 rcf for 5 minutes when they are suspended in 1-2 ml of buffer/medium. For larger cell suspension volumes, we recommend either 300 rcf for 10 minutes of 350 rcf for 5 minutes.

**Buffers**

The pH of buffers and percoll can decrease or increase over time. It is therefore important to make sure they are in the right range with a pH-meter the day of cell isolation, before starting the procedure.

*Adjusting the pH of percoll can be daunting. Store percoll at 4°C and adjust the pH of cold percoll. The pH of percoll is about 9, and this is harmful to the cells and leads to lower yield. In order to adjust the pH of percoll, we recommend pouring e.g. 80 ml of pure percoll into a 100 ml glass bottle, and place it onto a magnetic stirrer. Under stirring, apply 5M hydrochloric acid (HCl) at small steps. If adjusting 80 ml pure percoll, apply 5M HCl step-wise (starting with e.g. 50 μL) until pH 7.3-7.4. This bottle can be kept at 4°C and re-adjusted at each cell isolation. This pH adjusted percoll will be then mixed with e.g. 10x DPBS (Step 12).

**Age of the animals and specific tissues**

The protocol described above can be used to remove myelin and debris across all mouse ages from embryonic stages up to adulthood. Moreover, it has been successfully applied to dissociate different regions of the central nervous system, from spinal cord to cortical and subcortical areas.

**Challenging tissue**

This protocol is efficient for proper dissociation of adult mouse GL-261 gliomas. However, there may be other types of tumors which are harder to get dissociated. If that is the case, we recommend using enzyme mixes which work at 4°C (e.g. Accutase) in Hibernate medium. The tissue can be incubated in a sterile well-plate at 4°C (e.g. in a cold room) on an orbital shaker for e.g. 15-30 minutes. Thereafter, a finishing round with the Dounce homogenizer will complete the dissociation (Step 4). Of note: The Hibernate medium is suitable for maintenance of neural tissue in ambient $CO_2$, hence, there is no need for an incubator.

**Rat brain tissue/very large tissue pools**

When working with rat brain, if using dissected brain regions, e.g. hippocampi, from 1-2 adult rats, the ratios and volumes for 15 ml falcons can be used (see Table 1 above). If pooling larger amounts of brain tissue, e.g. hippocampi or cortices from more than 2 rats, we recommend the following modifications of the above protocol:

- Consider a Dounce homogenizer for larger volumes, e.g. 5 or 7 ml (with loose pestle!).
- Dissociate the tissue in 4 ml Hibernate in the Dounce homogenizer.
- Rinse the homogenizer and the cell-strainer 3-4 times.
- Step 11: pellet the dissociated tissue at 450 rcf for 10 minutes.
- Step 13: Resuspend the pellet in 4.5 ml DPBS. After resuspension, bring the volume to 6 ml with more DPBS (if necessary).
- Add 2 ml of isotonic percoll and mix well with a pipett-boy. (Final volume: 8ml).

- Delicately apply an overlay of 4 ml DPBS and centrifuge at 3000 rcf, 10 min.
- Proceed as above from Step 15.
